# Generalized Biological Foundation Model with Unified Nucleic Acid and Protein Language

**DOI:** 10.1101/2024.05.10.592927

**Authors:** Yong He, Pan Fang, Yongtao Shan, Yuanfei Pan, Yanhong Wei, Yichang Chen, Yihao Chen, Yi Liu, Zhenyu Zeng, Zhan Zhou, Feng Zhu, Edward C. Holmes, Jieping Ye, Jun Li, Yuelong Shu, Mang Shi, Zhaorong Li

## Abstract

The language of biology, encoded in DNA, RNA, and proteins, forms the foundation of life but remains challenging to decode due to its complexity. Traditional computational methods often struggle to integrate information across these molecules, limiting a comprehensive understanding of biological systems. Advances in Natural Language Processing (NLP) with pre-trained models offer new possibilities for interpreting biological language. Here, we introduce LucaOne, a pre-trained foundation model trained on nucleic acid and protein sequences from 169,861 species. Through large-scale data integration and semisupervised learning, LucaOne demonstrates an understanding of key biological principles, such as DNA-Protein translation. Using few-shot learning, it effectively comprehends the central dogma of molecular biology and performs competitively on tasks involving DNA, RNA, or protein inputs. Our results highlight the potential of unified foundation models to address complex biological questions, providing an adaptable framework for bioinformatics research and enhancing the interpretation of life’s complexity.

## Background

From the discovery of DNA to the sequencing of every living form, the faithful rule-based flow of biological sequence information from DNA to RNA and protein has been the central tenet of life science. These three major information-bearing biopolymers carry out most of the work in the cell and then determine the structure, function, and regulation of diverse living organisms[1, 2].

The basic information in the threes is presented in a linear order of letters: four nucleotides for DNA or RNA and 20 standard and several non-standard amino acids for proteins. Their secondary or higher structure also contains information attributed to biological functions and phenotypes. This genetic principle resembles the human linguistic system. Darwin wrote in his ***The Descent of Man***: ’The formation of different languages and distinct species, and the proofs that both have been developed through a gradual process, are curiously the same.’[3]. Various studies have testified to these parallels ever since, promoting the understanding and decoding of biological language[4–6].

Given the rapid advancements in machine learning technologies for human language processing, our efforts to decode biological language are bound to accelerate by leveraging insights from the former. The recent development of transformer architecture showed the superior capability of generalizing massive sequence-based knowledge from large-scale labeled and unlabeled data, which empowered language models and achieved unprecedented success in natural language processing (NLP) tasks. By pre-training on large datasets, foundational models learn the general characteristics of biological sequences. These models compute the input sequence into an embedding, a numerical representation that succinctly captures its semantic or functional properties. On this basis, various biological computation problems can be addressed through direct prediction, embedding analysis, or transfer learning[7]. In life science, substantial efforts have been put into adopting such language models, especially in protein tasks (ProTrans[8], ProteinBERT[9], ESM2[10], Ankh[11]), such as structure prediction[10, 12] and function annotation[13, 14]. In the realm of nucleic acid-focused tasks, several models have been introduced within niche areas (DNABert2[15], HyenaDNA[16], ScBert[17]). However, a broadly applicable, foundational model for nucleic acids remains elusive in widespread adoption across various disciplines.

Therefore, we have opted for a more fundamental and universal approach and developed a pre-trained, biological language semi-supervised foundation model, designated as ”LucaOne”, which integrates nucleic acid (DNA and RNA) and protein sequences for concurrent training. This methodology allows the model to process and analyze data from nucleic acids and proteins simultaneously, facilitating the extraction of complex patterns and relationships inherent in the processes of gene transcription and protein translation[18, 19].

We further examine that LucaOne exhibits an emergent understanding of the central dogma in molecular biology: the correlation between DNA sequences and their corresponding amino acid sequences, supporting the notion that the concurrent training of nucleic acid and protein sequences together yields valuable insights[20]. To illustrate LucaOne’s practical effectiveness, we present seven distinct bioinformatics computational scenarios. These examples highlight LucaOne’s ease of use in real-world applications and demonstrate its superior performance to state-of-the-art (SOTA) models and other existing pre-trained models.

## Results

### LucaOne as a Unified Nucleic Acid and Protein Foundation Model

LucaOne was designed as a biological language foundation model through extensive pretraining on massive datasets, enabling the extraction of generalizable features for effective adaptation to various downstream tasks, therefore allowing researchers to efficiently employ pre-trained embeddings from LucaOne for a diverse range of bioinformatics analysis, even when there is limited training data, thereby significantly enhancing their performance. This model leverages a multifaceted computational training strategy that simultaneously processes nucleic acids (DNA and RNA) and protein data from 169,861 species (only those with a minimum of 10 sequences within the training dataset are counted). Consequently, LucaOne possesses the capability to interpret biological signals and, as a foundation model, can be guided through input data prompts to perform a wide array of specialized tasks in biological computation.

**Fig. 1** depicts the LucaOne framework, which adopts and enhances the Transformer-Encoder[21] (**Methods A**). LucaOne’s vocabulary comprises 39 unique tokens representing nucleotides and amino acids (**Methods B**). We used pre-layer normalization to supersede post-layer normalization to make deep networks easier to train. Rotary Position Embedding (RoPE) replaces traditional absolute positional encoding for inferring longer sequences. Additionally, the mixed-training model distinguishes nucleotides and amino acids by utilizing token-type encoding, assigning 0 to nucleotides and 1 to amino acids. To comprehensively assimilate the patterns and structures pervasive in universal biological language and the inherent knowledge these patterns convey, we have compiled an extensive collection of nucleic acid and protein datasets as the foundational pre-training material. RefSeq provided nucleic acid sequences, including DNA and RNA, and annotations for eight selected Genome region types and their order-level taxonomy. Protein data included sequences (from UniProt and ColabFoldDB), annotations (from InterPro, UniProt, and ColabFoldDB), and tertiary structures (from RCSB-PDB and AlphaFold2) (**Fig. 2-a, Extended Data Figure 1, Extended Data Figure 2, and Supplementary** Figure 1). A semisupervised learning[19] approach was employed to enhance its applicability in biological language modeling. So, our pre-training tasks have been augmented with eight foundational sequence-based annotation categories. These annotations complement the fundamental self-supervised masking tasks, facilitating more effective learning for improved performance in downstream applications (**Fig. 2-b and Supplementary** Figure 3). Overall, LucaOne comprised 20 transformer-encoder blocks with an embedding dimension of 2,560 and a total of 1.8 billion parameters. The downstream task utilized a model checkpoint at 5.6M (**Methods E**). To illustrate the benefits of mixed training for nucleic acids and proteins, we trained the two additional models (LucaOne-Gene/LucaOne-Prot) separately using nucleic acids and proteins individually, and made a comparison using the same checkpoint in the central dogma of molecular biology task. Details of the pre-training data, pre-training tasks, and pre-training details refer to **Methods C, D, and E**, respectively.

**Fig. 1:**
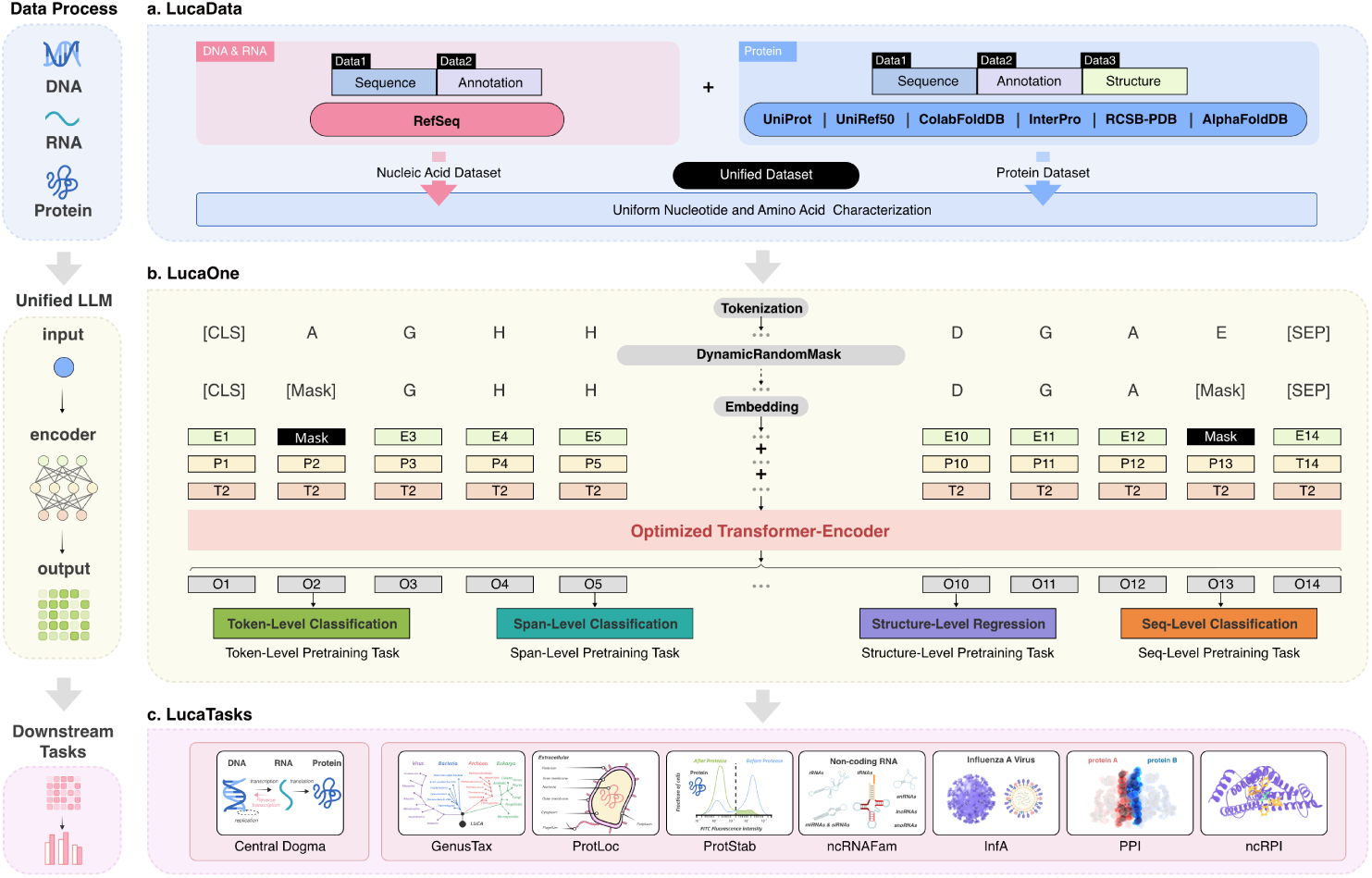
The workflow of LucaOne. **a.** Data source and processing for pre-training. The nucleic acid data was from RefSeq and included sequences and annotations, which consisted of order-level taxonomy and eight selected genome region types. Protein encompasses sequences (from Uniref50, UniProt, and ColabFoldDB-metagenomic protein collection (i.e. ColabFoldDB), where Uniref50 is clustered set of sequences from the UniProt with at least 50% sequence identity to enhance the learning of these representative sequences), annotations (order-level taxonomy from UniProt and ColabFoldDB, keywords from UniProt, and features such as sites, homologous superfamilies, and domains from InterPro), and tertiary structures (experimentally-determined structure from RCSB-PDB and predicted structure from AlphaFold2-Swiss-Prot). **b.** Pre-training model architecture and pre-training tasks. The Encoder is an improved transformer encoder. Based on two self-supervised mask tasks, an additional eight semi-supervised pre-training tasks were introduced to enhance the model’s understanding of the data through annotations in the sequences. **c.** Downstream tasks for validation based on LucaOne embedding. The representational capabilities of LucaOne were verified using eight downstream tasks, whose inputs include DNA, RNA, proteins, and their interrelated pairs.

**Fig. 2:**
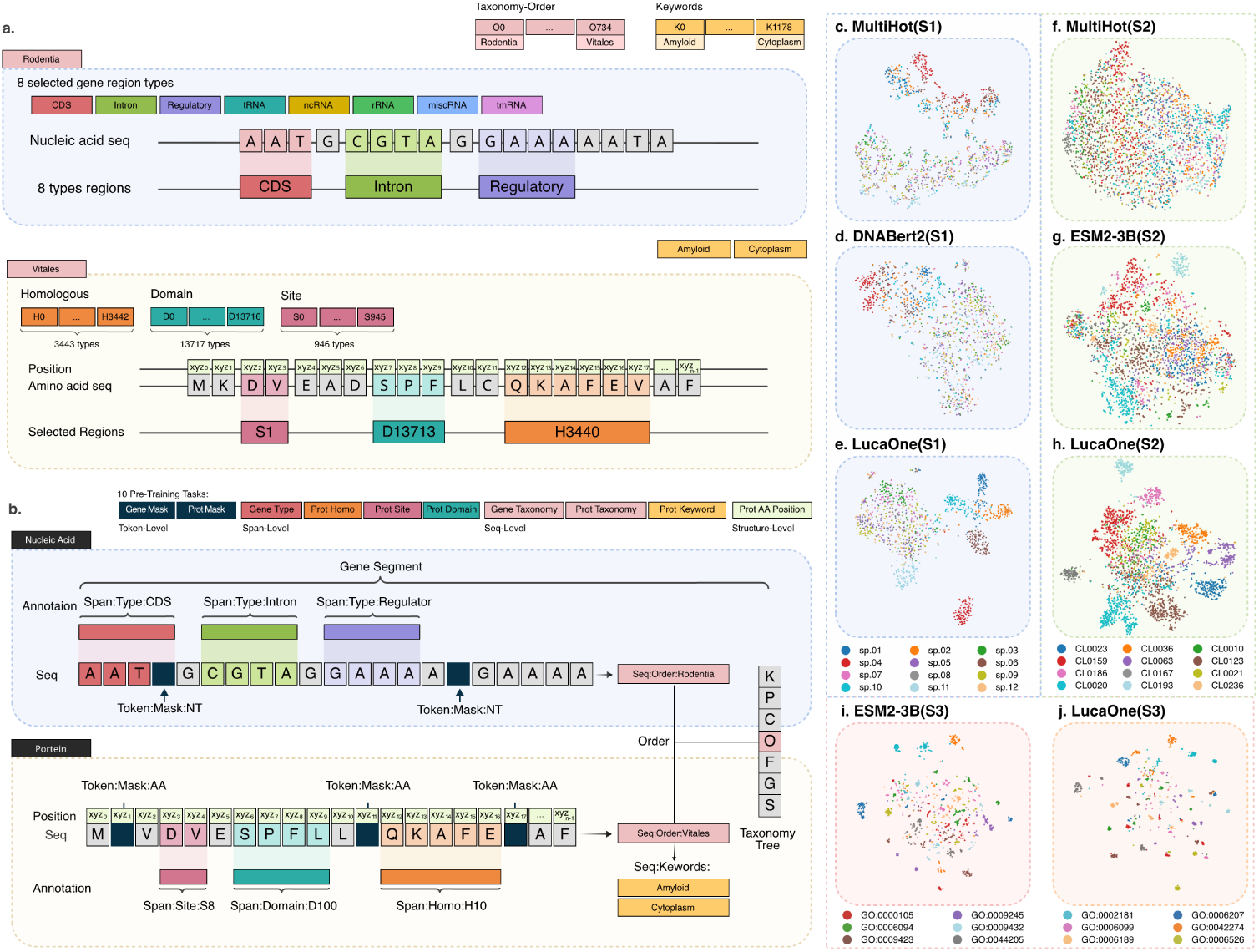
The data and tasks for pre-training LucaOne, and t-SNE on four embedding models. **a.** Details of pre-training data. Nucleic acids included sequence and two kinds of annotation. The protein consisted of sequence, five types of annotation, and tertiary structure coordinates. **b.** Details of pre-training tasks. The pre-training tasks included two self-supervised mask tasks and eight semi-supervised tasks. **c.**∼**j.** t-SNEs of the four embedding methods on the S1-nucleic acid contigs with 12 species from the CAMI2 database, S2-protein sequences across 12 clan categories from the Pfam database, and S3-protein sequences across the top 12 most prevalent Gene Ontology (GO) terms from the UniProt database. The results show that LucaOne’s representation has better clustering on these three datasets (nucleic acid sequences of the same species should be clustered because of high sequence similarity, and protein sequences of the same Pfam clan or GO term should be clustered of similar structures and functions). (**sp.01**: unclassified *Pseudomonas* species, **sp.02**: *Aeromonas salmonicida*, **sp.03**: unclassified *Vibrio* species, **sp.04**: *Streptomyces albus*, **sp.05**: *Aliivibrio salmonicida*, **sp.06**: unclassified *Brevundimonas* species, **sp.07**: *Vibrio anguillarum*, **sp.08**: *Aliivibrio wodanis*, **sp.09**: *Moritella viscosa*, **sp.10**: unclassified *Enterobacterales* species, **sp.11**: unclassified *Tenacibaculum* species, **sp.12**: unclassified *Aliivibrio* species; **GO:0000105**: L-histidine biosynthetic process, **GO:0009245**: lipid A biosynthetic process, **GO:0002181**: cytoplasmic translation, **GO:0006207**: ’de novo’ pyrimidine nucleobase biosynthetic process, **GO:0006094**: gluconeogenesis, **GO:0009432**: SOS response, **GO:0006099**: tricarboxylic acid cycle, **GO:0042274**: ribosomal small subunit biogenesis, **GO:0009423**: chorismate biosynthetic process, **GO:0044205**: ’de novo’ UMP biosynthetic process, **GO:0006189**: ’de novo’ IMP biosynthetic process, **GO:0006526**: L-arginine biosynthetic process).

We utilized t-distributed stochastic neighbor embedding (t-SNE) to visualize the embeddings from three distinct datasets: a nucleic acid dataset (S1), comprising sequences from 12 marine species, a protein dataset (S2), consisting of sequences from 12 clans (Pfam clans are groups of protein families that are evolutionarily related and share similar structures and functions.), and another protein dataset (S3), organizing of recently updated sequences from the top 12 most prevalent Gene Ontology (GO) terms-biological processes subset. This visualization was compared to the results obtained using the Multi-OneHot, DNABert2[15], and ESM2-3B[10] embedding approaches. The outcomes, as illustrated in **Fig. 2(c**∼**k)**, revealed that the embeddings produced by LucaOne were more densely clustered, indicating that this method may encapsulate additional contextual information beyond the primary sequence data. (Dataset S1, S2, and S3 details are in **Methods F**, and the embedding clustering metrics are in **Extended Data Table 1**). In addition, we examined the correlation between nucleic acid sequences and protein sequences of the same genes based on embeddings. The results demonstrated that, despite the absence of paired data and explicit correspondence relationships during training, the sequences (nucleic acids and proteins) of the same gene exhibited convergence within the LucaOne embedding space. Moreover, this convergence was more pronounced compared to other independently trained pre-trained models and sequence alignment methods (Details in **Methods F**).

### Learning Central Dogma of Molecular Biology

Our additional objective was to account for known gene and protein sequences occupying a minuscule yet biologically active niche within their respective design spaces, with a subset of these sequences exhibiting correspondence based on the central dogma. Consequently, throughout the training phase of the LucaOne model, we refrained from incorporating any explicit representations of the relationships between DNA, RNA, and protein sequence, seeking to test whether the model inherently grasped the correlation between the genetic and protein data[22, 23].

We designed an experimental task to assess the ability of LucaOne to recognize the inherent link between DNA sequences and their corresponding proteins. We have constructed a dataset comprising DNA and protein matching pairs derived from the NCBI-RefSeq database, with a proportion of 1: 2 between positive and negative samples (**Fig. 3-a, 3-b**, and **Methods G**). To better test whether the LucaOne model has already learned the correspondence between nucleic acid and protein sequences in the central dogma, few-shot learning was employed for validation. The samples were then randomly allocated across the training, validation, and testing sets in a ratio of 4: 3: 25, respectively (refer to ”Original Dataset” in the following sections).

**Fig. 3:**
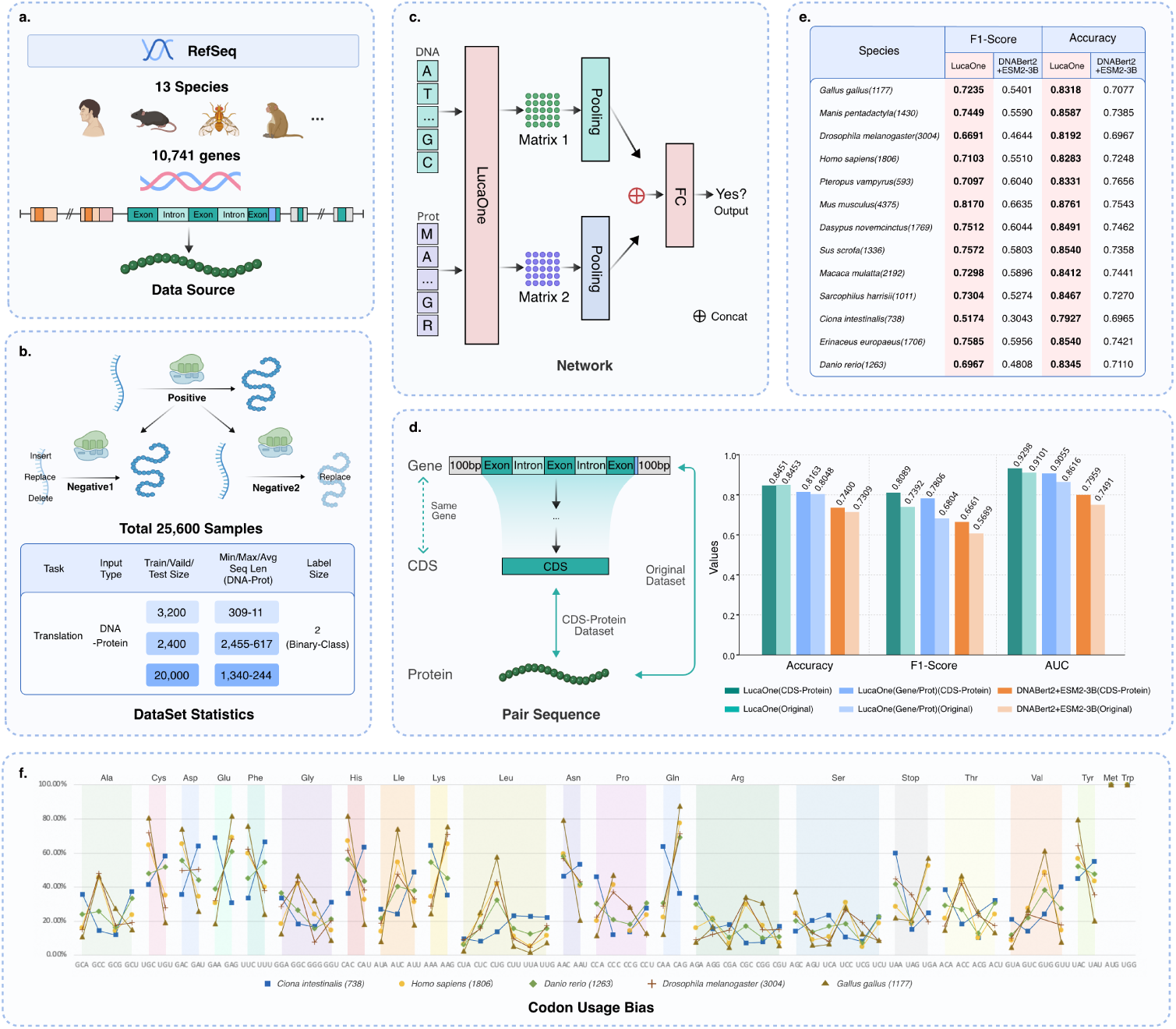
The workflow of the central dogma of molecular biology task. **a.** Dataset from 13 species with 10,471 genes in RefSeq. **b.** Prepared 8,533 positive samples and 17,067 negative samples and took a specific sample dividing strategy to test the model performance in this task (training set: validation set: testing set=4: 3: 25). **c.** Based on different embedding methods of DNA-Protein pair sequences, a simple downstream network was used for modeling and illustrating their representational ability. **d.** Models performance comparison (validation + testing Dataset) on original and CDS-Protein datasets. **e.** Comparative performance analysis (validation + testing Dataset) of the models across diverse species datasets (Sample counts in brackets). **f.** One species for each Class was selected to undergo a codon usage bias analysis, which adheres to the conventions of the standard genetic code; this entails comparing the relative usage frequencies of different codons for each amino acid, ensuring that the total adds up to 100%. The species *Ciona intestinalis* exhibits a codon usage bias that is markedly distinct from that of other species - overall lower GC content. Details in **Methods G**.

The study employed a simple downstream network to evaluate LucaOne’s predictive capacity (**Fig. 3-c**). LucaOne encoded nucleic acid and protein sequences into two distinct fixed embedding matrices (Frozen LucaOne). Then, each matrix was processed through pooling layers (either Max Pooling or Value-Level Attention Pooling[24]) to produce two separate vectors. The vectors were concatenated and passed through a dense layer for classification.

We compared the performance of different modeling approaches, including One-hot with a transformer, a transformer model with the random initialization, nucleic acid embeddings from DNABert2, protein embeddings from ESM2-3B, as well as two separate versions of the LucaOne foundation model trained independently on nucleic acid and protein sequences (LucaOne-Gene and LucaOne-Prot), and the unified training foundational version of LucaOne (**Fig. 3-d** and **Extended Data Table 2**). The findings indicated that modeling methods lacking pre-trained elements (One-hot and random initialization, see **Extended Data Table 2** were unable to acquire the capacity for DNA-protein translation in this dataset. In contrast, LucaOne’s embeddings were able to learn this capacity with limited training examples effectively and significantly surpassed both the amalgamation of the other two pretrained models (DNABert2 + ESM2-3B) and the combined independent nucleic acid and protein LucaOne models using the same dataset, architecture, and checkpoint. This suggests that pre-trained foundational models can provide additional information beyond the specific task samples for such biological computation tasks. Moreover, LucaOne’s unified training approach for nucleic acids and proteins enabled it to learn within a single framework, thereby capturing the fundamental intrinsic relationships between these two categories of biological macromolecules to some extent.

A CDS-Protein dataset using data from the original task was prepared to further evaluate the model’s capabilities. As **Fig. 3-d** shows that models trained exclusively on the CDS-Protein dataset demonstrated improvements across multiple performance metrics, including accuracy, F1-score, and AUC. When comparing the LucaOne model with the LucaOne-Gene/Prot model and the DNABert2 + ESM2-3B model, the enhancements were more substantial in the latter two model groups compared to LucaOne alone. It suggests that the LucaOne model possess marginally enhanced discriminative capabilities between coding and noncoding regions. However, our experimental results (**Supplementary** Figure 9) demonstrate a decline in LucaOne’s prediction accuracy as the number of exon within the target sequence region increases. This observed limitation represents a critical area for future model optimization. Furthermore, when evaluating performance across datasets from different species, both models show consistent results, except for a notable decrease in performance with *Ciona intestinalis*. This deviation can largely be attributed to its unique codon usage patterns, which differ significantly from other species in the study (**Fig. 3-e, 3-f**). Given the minimal sample size for this species in the dataset and with only 16% designated for training, it is likely that the models were unable to adequately learn the specific rules of the central dogma under these codon preferences, even though the analysis was conducted under the rule of the Standard Code. The observed divergence in codon preference suggests that *Ciona intestinalis* may possess more distinctive translation mechanisms from genetic material to proteins, which could be attributed to its unique evolutionary trajectory and selective pressures[25]. Furthermore, a dataset expanded with two urochordate species was utilized for model training and testing. The F1-score of the new model improved significantly for *Ciona intestinalis*, while the performance for other species remained comparable to that of the original model (**Methods G** and **Extended Data Table 3**). Based on this, it is inferred that with an expanded training data size encompassing a wider array of central dogma rules, LucaOne has the potential to more thoroughly assimilate the syntactical rules associated with genetic information processing, enabling its application to a more diverse set of scenarios.

### LucaOne Provides Embeddings for Diverse Biological Computational Tasks

To ascertain the capacity of the LucaOne model to provide effective embeddings for a variety of downstream tasks, we conducted validation studies across seven distinct downstream tasks, which include single-sequence tasks such as prediction of genus taxon (GenusTax), classification of ncRNA families (ncRNAFam), and the prediction of protein subcellular localization (ProtLoc) as well as the assessment of protein thermostability (ProtStab). For homogeneous paired-sequence tasks, we predicted influenza hemagglutination assays based on a pair of nucleic acid sequences (InfA) and assessed protein-protein interactions (PPI) utilizing pairs of protein sequences. Additionally, we forecasted the interactions between ncRNA and proteins (ncRPI) for the heterogeneous sequence task (Full task descriptions in **Methods H** and **Extended Data Table 4**).

For each task, we performed two types of comparative analysis: one against the state-of-the-art (SOTA) results and another using the same downstream network to assess LucaOne embeddings against the widely used nucleic acid and protein pre-trained language models, DNABert2 and ESM2-3B, respectively. These comparative analyses are instrumental in elucidating the incremental contributions of foundation models when addressing related analytical tasks and in evaluating the specific effectiveness of the embeddings generated by LucaOne with DNABert2 and ESM2-3B.

Similarly, we used a simple downstream network to facilitate processing these tasks. We illustrated the capacity of trained and frozen LucaOne to analyze nucleic acid (DNA and RNA) and protein sequences. **Fig. 4(a**∼**c)** displays the network architectures for three distinct input types. For tasks requiring paired inputs, a concatenation step is necessary to merge the output vectors of the pairs into a single extended vector. Finally, a fully connected (FC) layer was utilized for the ultimate output, which could be for classification or regression purposes. **Fig. 4(d**∼**k)** displays a comparative analysis of performance on seven distinct biomedical tasks, revealing that LucaOne demonstrates superior representational capabilities over competing models in the GenusTax, ProtStab, ncRNAFam, InfA, and PPI evaluations, and comparable performance on the other two: ProtLoc and ncRPI. Notably, within the nucleic acid-centric GenusTax and ncRNAFam, LucaOne’s accuracy has increased by 0.05 and 0.026, respectively, indicating a marked improvement over DNABert2. In the InfA task, LucaOne excelled with an exceptional accuracy of 1.0, reflecting its outstanding ability to represent this task data. For the ProtStab task, it surpassed ESM2-3B with a 0.015 increase in Spearman’s rank correlation coefficient (SRCC) and similarly showed a slight improvement in the evaluation of protein-protein interaction (PPI). Compared with DeepLocPro[26] in the task of ProtLoc, LucaOne was competitive with ESM2-3B and demonstrated a 0.025 accuracy improvement. Although LucaOne did not outperform the elaborate network model ncRPI-LGAT[27] in evaluating ncRPI, it still exceeded the combined abilities of DNABert2 and ESM2-3B. LucaOne’s effectiveness was particularly notable in processing tasks involving heterogeneous sequences of nucleic acids and proteins; employing a unified representation model is advantageous compared to using separate models. The outcomes of these tasks underscored the robust representational capabilities of LucaOne for both nucleic acid and protein sequences. LucaOne could improve performance across a spectrum of downstream tasks, streamline networks for downstream tasks, and reduce computational resource demands (More results of hyperparameter comparison experiments and detailed metrics in **Methods I**, **Extended Data Table 5**, and Supplementary Figure 4).

**Fig. 4:**
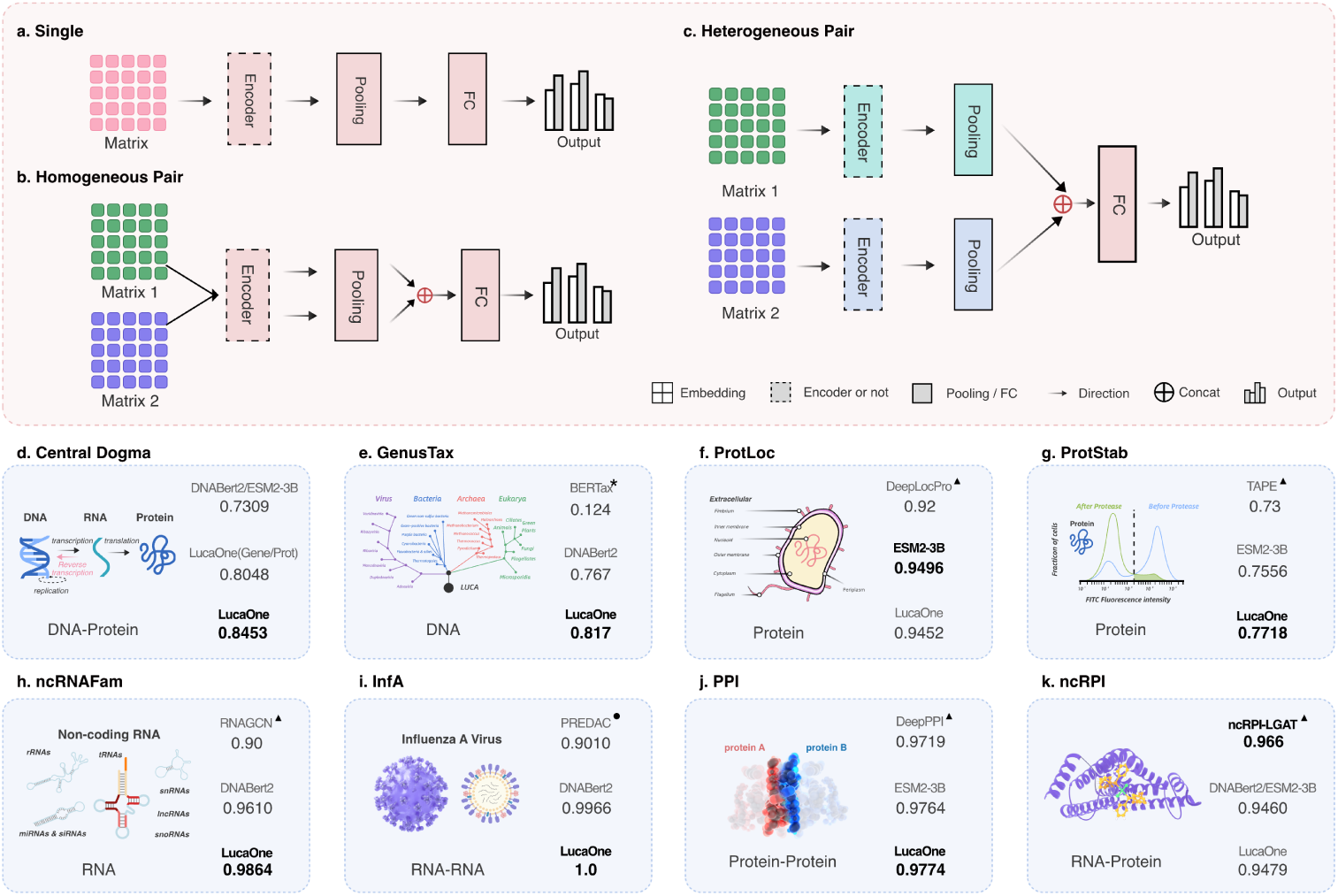
Downstream task networks with three input types and results comparison of 8 verification tasks. Based on the embedding matrix, three types of inputs in the downstream task are corresponding networks: **a.** A single sequence, including GenusTax, ncRNAFam, ProtLoc, and ProtStab; **b.** Two same-type sequences, including InfA and PPI; **c.** Two heterogeneous sequences: Central Dogma and ncRPI. **d.**∼**k.** Comparison results of 8 downstream tasks. The Spearman Correlation Coefficient (SRCC) was employed for the ProtStab regression task, and Accuracy (Acc) was used for other tasks. Comparative methods include the SOTA, DNABert2-based (for nucleic acids), ESM2-3B-based (for proteins), and LucaOne-based. The top right ★ indicates inference using the trained method, the top right ▴ indicates direct use of the results in its paper, and the top right • indicates repetition using its method and is better than the results in the paper.

## Discussion

The attempt to build a universal biological language model is to develop a sophisticated cataloging and retrieval system for ”The Library of Mendel” - the genetic version of ”The Library of Babel.”[28, 29]. The diversity of genetic variations presents a vast ”design space” that is arguably as rich as the entirety of human literature, if not more so, given the far longer history of life on Earth compared to our record of literature. However, in stark contrast, the proportion of genetic sequences we have successfully identified and cataloged remains significantly smaller than the volume of documented human languages. Moreover, the growth of our understanding and documentation of this ’biological language’ is unlikely to occur suddenly or rapidly[30, 31]. Our endeavor herein offers a computational model that posits the potential to represent the paradigm of biological language. However, we must temper our expectations regarding this model’s rapid and seamless refinement toward an idealized state of perfection. In developing the LucaOne model, we used deep learning frameworks and techniques from natural language processing. However, we observed systemic discrepancies when applying these models, which were highly successful in natural language contexts, to genomic language[32]. The architecture of BERT-based pre-trained language models focuses on understanding context but may not efficiently capture biological sequences’ unique attributes and characteristics[33, 34]. Furthermore, the functions and expressions of biological sequences are not solely determined by their genetic sequences but also by the environment in which they are expressed - a factor for which there is presently no practical modeling approach. Standardized methods for processing annotated or phenotypic data are lacking, which can lead to inaccuracies and omissions[35, 36]. Moreover, the continual learning and scalability aspects have yet to be fully explored in this study, primarily due to resource constraints. As a result, the complexities of the model’s learning capabilities have not been thoroughly examined at this point, highlighting the primary area of research for the subsequent phase[37]. In terms of application, due to the diversity of contexts, a robust evaluation system is absent for generalizability and domain adaptability, with small, specialized models occasionally outperforming large pre-trained models in conjunction with downstream tasks in certain areas[32, 38].

In light of these considerations, researchers may need to develop specialized pre-trained models tailored to genomic language to improve encoding and comprehension of biological data, ensuring adaptability across a broader spectrum of computational biology tasks. Promising directions include architectural innovations in pre-training models, such as incorporating genetic programming concepts into large language models (LLMs)[39, 40]. Another avenue is to harmonize multimodal data, encompassing sequences, feature annotations, experimental results, images, and phenotypical information to better understand biological systems beyond unsupervised sequence data learning[41, 42]. Additionally, employing more transparent algorithms may enhance the interpretability of the model, facilitating better integration with existing biological research frameworks and model development[43, 44]. Lastly, given the necessity for pre-trained models to efficiently fine-tune or apply to downstream tasks, paradigms need to expedite model adaptation to new tasks and broader application contexts[32].

To conclude, this paper documented our effort to build a comprehensive large model to represent the intricacies of the biological world. The capabilities demonstrated by LucaOne showed considerable promise and highlighted several areas that necessitate substantial advancements. Such multimodal pre-trained foundational models, grounded in bioinformatics, will prove immensely valuable in accelerating and enhancing our comprehension of biological phenomena.

## Methods

### **A.** Model Architecture

**Fig. 1-b** illustrates the design of LucaOne, which utilizes the Transformer-Encoder[21] architecture with the following enhancements:

1. The vocabulary of LucaOne comprises 39 tokens, including both nucleotide and amino acid symbols (refer to **Methods B**);
2. The model employs Pre-Layer Normalization over Post-Layer Normalization, facilitating the training of deeper networks[45];
3. Rotary Position Embedding (RoPE[46]) is implemented instead of absolute positional encoding, enabling the model to handle sequences longer than those seen during training;
4. It incorporates mixed training of nucleic acid and protein sequences by introducing token-type embeddings, assigning 0 for nucleotides and 1 for amino acids;
5. Besides the pre-training masking tasks for nucleic acid and protein sequences, eight semi-supervised pre-training tasks have been implemented based on selected annotation information (refer to **Methods D**).

### **B.** Vocabulary

The vocabulary of LucaOne consists of 39 tokens. Due to the unified training of nucleic acid and protein sequences, the vocabulary includes four nucleotides (’A’, ’T’, ’C’, ’G’) of nucleic acid (’U’ compiled with ’T’ in RNA), ’N’ for unknown nucleotides, 20 amino acids of protein (20 uppercase letters excluding ’B’, ’J’, ’O’, ’U’, ’X’, and ’Z’), ’X’ for unknown amino acids, ’O’ for pyrrolysine, ’U’ for selenocysteine, other three letters (’B’, ’J’, and ’Z’) not used by amino acids, five special tokens: ’[PAD]’, ’[UNK]’, ’[CLS]’, ’[SEP]’, ’[MASK]’, and three other characters: (’.’, ’-’, and ’*’). Due to the amino acid letters overlapping with the nucleotide letters, the use of ’1’, ’2’, ’3’, ’4’, and ’5’ instead of ’A’, ’T’, ’C’, ’G’, and ’N’, respectively.

### **C.** Pre-training Data Details

#### Nucleic Acid

The nucleic acid was collected from the NCBI RefSeq genome database, involving 297,780 assembly accessions. The molecular types included DNA and RNA (**Fig. 2-a**). The DNA sequence, DNA selected annotation, RNA sequence, and RNA selected annotation were obtained from the format files ’genomic.fna’, ’genomic.gbff’, ’rna.gbff’, and ’rna.fna’, respectively. Among all pre-training sequences, 70% of DNA sequences and 100% of RNA sequences were derived from annotated genomes, while the remaining unannotated sequences were retained to ensure diversity.

**DNA reverse strand:** The DNA reverse strand also contains a lot of annotation information, so the DNA dataset expanded reverse strand sequences with their annotation. A total of 23,095,687 reverse-strand DNA sequences were included.

**Genome region types:** Eight important genome region types in nucleic acids were selected, including ’CDS’, ’intron’, ’tRNA’, ’ncRNA’, ’rRNA’, ’miscRNA’, ’tmRNA’, and ’regulatory’. Each nucleotide in the sequence had a label index of 8 categories (0∼7) or *-100* when it did not belong to these eight categories.

**Order-level taxonomy:** The order-level label of the taxonomy tree was selected as the classification label of the nucleic acid sequence. Each sequence had a label index of 735 categories (0∼734) or *-100* without the order-level taxonomy.

**Segmentation:** Due to the limited computing resources, each nucleic acid sequence was segmented according to a given maximum length. The fragmentation strategy was presented in **Supplementary** Figure 2.

#### Protein

Protein sequence data were obtained from UniRef50, UniProt, and ColabFoldDB. UniRef50 was added to the UniProt database to upsampling high-quality representative sequences, while ColabFoldDB was incorporated to enhance the diversity of protein sequences. For ColabFoldDB, redundancy within each cluster was minimized by retaining only the ten most diverse sequences. Duplicated sequences between UniProt and ColabFoldDB were excluded. As shown in **Fig. 2-a**, compared to nucleic acids, proteins contained more information, including sequence, taxonomy, keywords, sites, homology regions, domains, and tertiary structure. Sequence, taxonomy, and keywords were collected from UniProt and ColabFoldDB. The sites, domains, and homology regions were extracted from Interpro. The tertiary structure was derived from RCSB-PDB and AlphaFold2-Swiss-Prot.

**Sequence:** The *right truncation* strategy was applied when the sequence exceeded the maximum length.

**Order-level taxonomy:** Order-level classification information is used as the protein sequence taxonomy. There were 2,196 categories; each sequence had a label index (0∼2,195) or *-100* if its order-level information was missing.

**Site:** Four types of site regions (‘Active site’, ‘Binding site’, ‘Conserved site’, and ‘PTM’) with 946 categories were included. For each amino acid in a sequence, if it was a site location, there was a label index (0∼945); otherwise, it was marked with *-100*.

**Homology:** A homologous superfamily is a group of proteins that share a common evolutionary origin with a sequence region, reflected by similarity in their structure. There were 3,442 homologous region types; Each amino acid in these regions had a label index (0∼3,441) corresponding to its type, and the other amino acids were labeled *-100*.

**Domain:** Domain regions are distinct functional, structural, or sequence units that may exist in various biological contexts. A total of 13,717 domain categories were included; Each amino acid in these regions had a label index (0∼13,716) corresponding to its category, and the other amino acids were marked with *-100*.

**Keyword:** Keywords are generated based on functional, structural, or other protein categories. Each sequence was labeled as a set of label indices with 1,179 keywords or *-100* without keywords.

**Structure:** The spatial coordinates of the *C_α_-atom* were used here as the amino acid coordinates. Each amino acid was labeled with a three-dimensional coordinate normalized within the protein chain. The amino acids at the missing locations were labeled *(−100, −100, −100)*. We obtained the experimentally determined structure in RCSB-PDB and the predicted structure by AlphaFold2 of UniProt (Swiss-Prot) and preferentially selected the structure in RCSB-PDB. In total, only about half a million protein sequences had structural information.

### **D.** Pre-training Tasks Details

LucaOne has employed a semi-supervised learning approach to enhance its applicability in biological language modeling. Unlike the traditional natural language machine learning domain, where tasks involve input and output of the same textual modality, bioinformatics analysis often involves different modalities for input and output data. Most bioinformatics downstream tasks extend from understanding nucleic acid or protein sequences, so our pre-training tasks have been augmented with eight foundational sequence-based annotation categories. These annotations complement the fundamental self-supervised masking tasks, facilitating more effective learning for improved performance in downstream applications. The selection criteria for these annotations focused on universality, lightweight design, and high confidence level; consequently, only a subset of the data possess such annotations. As listed in **Supplementary** Figure 3, there are 10 specific pre-training tasks at four levels.

**Token Level Tasks:** Gene Mask and Prot Mask tasks randomly mask nucleotides or amino acids in the sequence following the BERT masking scheme [47] and predict these masked nucleotides or amino acids based on the sequence context in training. The loss functions of the two mask language pretraining tasks are defined in **Supplementary Notes 1 Eq. (1)**∼**(2)**.

**Span Level Tasks:** The model is trained to recognize some essential regions based on the sequence context. For nucleic acid sequences, eight essential genome region types are learned. For protein sequences, three types of regions are labeled: site, homology, and domain regions. The loss functions of these span-level pertaining tasks are listed in **Supplementary Notes 1 Eq. (3)**∼**(7)**.

**Seq Level Tasks:** Gene-taxonomy, Prot-taxonomy, and Prot-keyword are the order-level taxonomies of nucleic acid, protein, and protein-tagged keywords, respectively. They are all sequence-level learning tasks. The loss functions of the three sequence-level pre-training tasks are presented in **Supplementary Notes 1 Eq. (8)**∼**(9)**.

**Structure Level Tasks:** Since the structure of a protein determines its function, we use a small amount of protein data with a tertiary structure for simple learning in the pre-training phase. Instead of learning the spatial position at the atomic level, the spatial position of amino acids is trained (using the position of the *C_α_ atom* as the position of the amino acid). The loss function of structure-level pertaining task is defined in **Supplementary Notes 1 Eq. (10)**.

**Data Processing:** All data processing tasks were performed on the Alibaba Cloud MaxCompute platform.

### **E.** Pre-training Information

On the dimensions of the embedding, the research conducted by Elnaggar et al. [11] demonstrates that the ESM2-3B model, with an embedding dimension of 2,560, significantly enhances performance compared to its counterpart, ESM2-650M, harbors an embedding dimension of 1,280. However, when the parameter count increases to 15 billion with an embedding dimension of 5120, there is no significant increase in performance. It is also observed that the ESM2-15B model requires substantially more time than ESM2-3B and ESM2-650M regarding the relationship between input sequence length and computational time. For the relationship between model size and training data size, Hoffmann et al. suggest that a minimum of 20.2 billion tokens is essential to adequately train a 1B-sized model [48].

Taking into account these insights, along with the consideration of available computational resources and the volume of pre-training data, some of the critical hyperparameters we adopted are as follows: the architecture of LucaOne consists of 20 transformer-encoder blocks with 40 attention heads each, supports a maximal sequence length of 1,280, and features an embedding dimension of 2,560. The model is composed of a total of 1.8 billion parameters. We employed 10 different pretraining tasks, assigning an equal weight of 1.0 to Gene-Mask, Prot-Mask, Prot-Keyword, and Prot-Structure tasks, while assigning a reduced weight of 0.2 to the remaining tasks to equilibrate task complexity (**Supplementary Notes 1 Eq. (11)**). We used the AdamW optimizer with betas (0.9, 0.98) and a maximum learning rate of 2e-4, incorporating a linear warm-up schedule throughout the learning rate updates. For the model training regimen, we utilized a batch size of 8 coupled with a gradient accumulation step of 32. The model underwent training on 8 Nvidia A100 GPUs spanning 120 days. A model checkpoint of 5.6 million (5.6M, trained with 36.95B tokens) was selected for the subsequent validation tasks, aligned with ESM2-3B in terms of the volume of data trained for comparison.

To elucidate the advantages of mixed training involving both nucleic acids and proteins, we further conducted experiments with two supplementary models, LucaOne-Gene and LucaOne-Prot, trained exclusively with nucleic acids and proteins, respectively. Their performance in the central dogma of the biology task was evaluated with the same checkpoint (5.6M) of the two models.

**Checkpoint Selection Criteria:** We have released the 5.6M checkpoint aligned with the ESM2-3B model for a comparable volume of data trained, which was trained with 36.95 billion tokens over 20 times the model’s parameters. We also released the 17.6M checkpoint (trained with 116.62B tokens) based on three criteria: **1)** The loss curve slowly descended after 17.6M steps during training (**Extended Data** Figure 3-a); **2)** The losses are relatively stable on the validation and testing set between 15M and 20M steps, making 17.6M optimal (**Extended Data** Figure 3-b **and 3-c**); **3)** The improvement in the performance of representative downstream tasks is very limited. For example, in the ncRPI task, the accuracy is 94.93% at checkpoint 17.6M, which is only a marginal improvement compared to an accuracy of 94.78% at checkpoint 5.6M (**Extended Data** Figure 3-d).

### **F.** LucaOne Embeddings Level Analysis

**Details of t-SNE Datasets**: The S1 dataset was curated from marine data available in CAMI2[49], selecting contigs with lengths ranging from 300 to 1,500 nucleotides. We focused on species that were identifiable and possessed at least 200 contigs. The contigs of each species were redundant by MMSeqs, employing a coverage threshold of 80% and sequence identity of 95%, culminating in a collection of 37,895 nucleic acid contigs from 12 species. We randomly selected 100 samples from each species, totaling 1,200 items for visualization.

The S2 dataset originated from clan data within Pfam, maintaining clan categories with a minimum of 100 Pfam entries, resulting in 189,881 protein sequences across 12 clan categories. For visualization, we randomly selected one sample for each Pfam entry in every clan, amounting to 2,738 samples.

The S3 dataset was selected from the UniProt database from May 1, 2023, to December 16, 2024, which does not overlap the pre-training data of LucaOne (before May 29, 2022). This data set focused on the lowest-tier Gene Ontology (GO) annotations within the hierarchical annotation framework of the biological processes (BP) subset, identifying the 12 most prevalent terms at this foundational level. Each GO term randomly samples 100 sequences between 100 and 2,500 amino acids in length, resulting in 1,200 protein sequences across the 12 GO terms (**Supplementary Note 2**).

**Convergence of Nucleic Acid and Protein Sequences for the Same Gene:** We prepared an additional dataset comprising nucleic acid and protein sequences for the same genes and examined their correlations solely on embeddings. The results indicated that, despite nucleic acid and protein sequences not being paired during model training, those corresponding to the same gene demonstrated convergence within the LucaOne Embedding Space. More details in **Supplementary Note 6** and **Supplementary** Figure 12.

**Task on Codon Degeneracy:** We designed an additional task based on influenza virus HA sequence data to verify whether LucaOne can distinguish between synonymous and non-synonymous mutations in a zero-shot manner, more details in **Supplementary** Figure 16.

**Task on Pseudogene Correction**: We conducted a mask task prediction analysis (zero-shot) on the data of the true gene (protein coding) and pseudogene pairs, the higher pseudogene correction rate and the true gene recovery rate demonstrated the model’s ability to identify the differences between pseudogenes and functional genes. More details in **Supplementary Note 7**, **Supplementary** Figure 13, and **Supplementary** Figure 14.

### **G.** Details of Central Dogma Tasks

**Dataset Construction-Original Dataset:** We devised an experimental task to determine whether LucaOne has established the intrinsic association between DNA sequences and their corresponding proteins. A total of 8,533 accurate DNA-protein pairs from 13 species were selected in the NCBI-RefSeq database, each DNA sequence extending to include an additional 100 nucleotides in the 5’ and 3’ contexts, preserving intron sequences within the data. In contrast, we generated double the number of negative samples by implementing substitutions, insertions, and deletions within the DNA sequences or altering amino acids in the protein sequences to ensure the resultant DNA sequences could not be accurately translated into their respective proteins, resulting in a total of 25,600 samples - DNA-protein pairs. Then the positive and negative samples were each subjected to random shuffles and subsequently divided into 32 equally sized subsets. Then these subsets were assigned to the training, validation, and testing sets in a 4: 3: 25 ratio. For more details, then see **Extended Data Table 4** and **Data Availability**

**Analysis of Misclassified Samples:** We analyze the misidentified samples from two perspectives: sequence and embedding. The relationship between sequence identity metrics and the prediction accuracy of the LucaOne embedding was presented in **Extended Data** Figure 4-a and **4-b**. Data details were presented in **Supplementary Note 3**. **Extended Data** Figure 4-a and **4-b** show that the prediction accuracy of LucaOne for mutated sample pairs improved as sequence similarity decreased. We also evaluated the embedding distance alterations corresponding to modifications in nucleic acid and protein sequences by employing mean pooling to calculate these distances. As illustrated in **Extended Data** Figure 4-c **and 4-d**, greater changes in embedding distances were correlated with improved predictive precision.

**Dataset Construction-2 More Species of Urochordates:** We incorporated two species with annotated reference genome urochordates (referred to as Tunicate in the NCBI Taxonomy) into our dataset: *Styela clava* (ASM1312258v2, GCF 013122585.1) and *Oikopleura dioica* (OKI2018 I68 1.0, GCA 907165135.1). For each of these urochordate species, 480 genes were randomly selected, and positive gene samples, nucleic acid negative samples, and protein negative samples were constructed using the same approach as in the original dataset. The same data shuffling and partitioning principles were applied and integrated with the original dataset to retrain the Central Dogma model. Data details and model performance are presented in **Extended Data Table 3**, **Extended Data** Figure 5, and **Data Availability**.

**Comparative Performance Analysis:** Upon integrating two additional urochordate species data, Dataset Version 2 as the model exhibited performance comparable to the original dataset models across all species except *Ciona intestinalis*. In particular, the F1-score for *Ciona intestinalis* improved significantly, despite the nearly unchanged accuracy. These findings suggest that supplementing the dataset with species that utilize a codon code similar to *Ciona intestinalis* enhances the model’s sensitivity to DNA-Protein correlations in these organisms while preserving its sensitivity to DNA-Protein correlations in species adhering to the standard codon code. For more details, please see **Extended Data Table 3** and **Data Availability**.

**CDS-Protein Task:** In the current NCBI RefSeq database, genomes with complete intron annotations are limited, and the accuracy of intron predictions from alternative tools may directly impact model performance. To mitigate these challenges, coding sequence (CDS) regions corresponding to genes in the original dataset were extracted as intron-free nucleic acid sequences to perform the same task. See **Supplementary Note 4** for data details and **Fig. 3-d** for analysis.

**Task for Cross-Species Homologous Gene Pairs:** We designed an additional task related to the central dogma by modifying the negative samples in the original study. Instead of manually altering the sequences, the negative samples were replaced with homologous genes from closely related species. Please refer to **Supplementary Note 5** for details.

### **H.** Downstream Tasks Details

**Genus Taxonomy Annotation (GenusTax):** This task is to predict which genus (taxonomy) the nucleic acid fragment may come from, which is very important in metagenomic analysis. A comparative dataset was constructed utilizing NCBI RefSeq, comprising 10,000 nucleic acid sequences, each extending 1,500 nucleotides and annotated with labels corresponding to 157 distinct genera (distributed as 33, 50, 29, and 45 across the four kingdoms: Archaea, Bacteria, Eukaryota, and Viruses, respectively). The dataset was randomly segregated into training, validation, and testing sets, adhering to an 8: 1: 1 partitioning ratio. It is important to note that while the LucaOne pre-training task utilized taxonomy annotations at the order level, the current task employs more granular genus-level annotations, thereby preventing label information contamination. This investigation constitutes a multi-class classification challenge. This dataset was also employed for two additional analyses: predicting the taxonomy of sequences at SuperKingdom and Species levels. The details are presented in **Extended Data Table 5**.

**Prokaryotic Protein Subcellular Location (ProtLoc):** This task is to predict the subcellular localization of proteins within prokaryotic cells, which has garnered substantial attention in proteomics due to its critical role[50]. It involves classifying proteins into one of six subcellular compartments: the cytoplasm, cytoplasmic membrane, periplasm, outer membrane, cell wall and surface, and extracellular space. Our approach adopted the same dataset partitioning strategy as DeeplocPro[26], a model based on experimentally verified data from the UniProt and PSORTdb databases. Our research undertakes a multi-class classification challenge, categorizing proteins based on their distinct subcellular localizations. For this dataset, we additionally designed a task based on the corresponding nucleic acid embeddings of the proteins. The result showed that embeddings derived from nucleic acid sequences are applicable to the task related to their corresponding protein sequences. The dataset and analytical results are provided in **Supplementary Note 8**.

**Protein Stability (ProtStab):** The evaluation of protein stability is paramount for elucidating the structural and functional characteristics of proteins, which aids in revealing the mechanisms through which proteins maintain their functionality in vivo and the circumstances that predispose them to denaturation or deleterious aggregation. We utilized the same dataset from TAPE[51], which includes a range of denovo-designed proteins, natural proteins, mutants, and their respective stability measurements. It is a regression task; each protein input

1. (*x*) correlates with a numerical label(*y* ∈ ℜ), quantifying the protein’s intrinsic stability.

**Non-coding RNA Family(ncRNAFam):** Non-coding RNA(ncRNA) represents gene sequences that do not code for proteins but have significant functional and biological roles. The objective is to assign ncRNA sequences to their respective families based on their characteristics. For this purpose, we utilize the dataset from the nRC[52], which is consistent with the data employed in the RNAGCN[53] study. Our methodology adheres to the same data partitioning into training, validation, and testing sets as done in these previous studies, enabling direct comparison of results. This project involves a multi-class classification challenge that encompasses 88 distinct categories.

**Influenza A Antigenic Relationship Prediction (InfA):** One of the foremost tasks in influenza vaccine strain selection is monitoring Hemagglutinin (HA) variant emergence, which induces changes in the virus’s antigenicity. Precisely predicting antigenic responses to novel influenza strains is crucial for developing effective vaccines and preventing outbreaks. The study utilizes data from the PREDAC[54] project to inform vaccine strain recommendations. Each data pair in this study comprises two RNA sequences of the HA fragment from distinct influenza strains, accompanied by corresponding antigenic relationship data. The objective is framed as a binary classification task identifying the antigenic similarity or difference between virus pairs.

**Protein-Protein Interaction (PPI):** The forecasting of protein-protein interaction networks represents a significant area of research interest. Our study utilized the DeepPPI[55] database, whose positive dataset samples were sourced from the Human Protein Reference Database after excluding redundant interactions, leaving 36,630 unique pairs. This dataset was randomly partitioned into three subsets: training (80%), validation (10%), and testing (10%). The primary objective of this research is to perform binary classification of protein-protein interaction sequences.

**ncRNA-Protein Interactions (ncRPI):** An increasing number of functional non-coding RNAs (ncRNAs), such as snRNAs, snoRNAs, miRNAs, and lncRNAs, have been discovered. ncRNAs play a crucial role in many biological processes. Experimentally identifying ncRNA-protein interactions (ncRPI) is typically expensive and time-consuming. Consequently, numerous computational methods have been developed as alternative approaches. For comparison, we have utilized the same dataset as the currently best-performing study, ncRPI-LGAT[27]. It is a binary classification task involving pairs of sequences.

### I. Comparison Result Detaills

We conducted a series of comparative experiments. According to **Fig. 4**, for all embedding methods, we compare whether the transformer encoder and two pooling strategies (max pooling and value-level attention pooling) were used on the model. At the hyperparameter level, we compared the number of encoder layers with the number of heads (4 layers with 8 heads and 2 layers with 4 heads), the peak learning rate of the Warmup strategy (1e-4 and 2e-4), and the batch size (8 and 16). **Extended Data Table 5** shows the result of comparing whether the encoder was used and which pooling method was used accordingly, and **Supplementary** Figure 4 shows more metrics on comparison experiments.

In the **ProtLoc** task, LucaOne’s accuracy is very close to that of the ESM2-3B; In the **ncRPI** task, the accuracy of the simple network with LucaOne’s embedding matrix is less than that of ncRPI-LGAT[27] but higher than that of DNABert2 + ESM2-3B; In **the other five tasks**, we achieved the best results. It is better not to use an encoder for **ProtLoc**, **InfA**, **PPI**, and **ncRPI** tasks. Using the Max Pooling strategy straightforwardly for the **ncRNAFam** and **GenusTax** tasks can obtain better results. We extended two tasks, four superkingdoms, and 180 species prediction tasks for the genus classification task with the same sequence data. LucaOne’s accuracy improved by 0.1 and 0.054, respectively. In particular, LucaOne is more effective than other large models in embedding sequences without an encoder.

## Data Availability

The pre-training dataset of LucaOne is opened at: http://47.93.21.181/lucaone/PreTrainingDataset/. The datasets of all downstream tasks are available at[56]: https://doi.org/10.5281/zenodo.15171943. Other supplementary materials are available at[56]: https://doi.org/10.5281/zenodo.15171943.

## Code Availability

The LucaOne’s model code is available at: https://github.com/LucaOne/LucaOne. The trained checkpoint files are available at[56]: https://doi.org/10.5281/zenodo.15171943. LucaOne’s embedding inference code is available at: https://github.com/LucaOne/LucaOneApp. The project of all downstream tasks is available at: https://github.com/LucaOne/LucaOneTasks. The trained checkpoint files of all downstream tasks are available at[56]: https://doi.org/10.5281/zenodo.15171943.

## Supporting information

Supplementary Datasheet

## Acknowledgments

This work was supported by the National Natural Science Foundation of China (82341118). M. S. is funded by the Shenzhen Science and Technology Program (KQTD20200820145822023), the Guangdong Province ”Pearl River Talent Plan” Innovation and Entrepreneurship Team project (2019ZT08Y464), the Guangzhou National Laboratory Major Project (GZNL2023A01001). Y.-F. P. is funded by National Natural Science Foundation of China (NSFC) Basic Research Project for Doctoral Students (Grant No.323B2018).

## Author Contributions

Conceptualization: Y. H., Z.-R. L., and M. S.; Model development and data preparation for LucaOne: Y. H., Y.-H. W., Y.-F. P., and Y.-C. C.; Downstream tasks understanding and models training: Y. H., P. F., Y.-T. S., and Y.-H. C.; Original draft: Y. H., Z.-R. L., and P. F.; Writing - Review and Editing: All authors; Graphic presentation design: Y. L. and Y. H.; Engineering leadership and resource acquisition: Z.-Y. Z. and J.-P. Y.; Science leadership and resource acquisition: J. L., E.-C. H., Z. Z., F. Z., and Y.-L. S.; Supervision: Y. H., M. S., and Z.-R. L.. We thank J. Wang, D.-C. Ma, and D.-Z. Shi for computational resource coordination. We thank H.-W. Zhang for maintaining computational resources and optimizing specific computing tasks at Yunqi Academy of Engineering (Hangzhou). We thank Y.-Q. Liu and M. Zhou for their participation in the subsequent technical validation in conjunction with this research. We thank X.-J. Du, W.-C. Wu, J.-Y. Yang, and S.-Q. Mei from Sun Yat-sen University (Shenzhen) for a valuable conversation on the development of the method, especially on understanding the downstream tasks. Last but not least, we thank Charles Darwin, Richard Dawkins, Steven Pinker, and Daniel Dennett for their profound insights that led to the early conceptual foundations of this study and guided its development pathway.

## Computational Resources

The data processing and training operations for LucaOne were carried out on Alibaba Cloud Computing Co., Ltd. Additionally, several tasks related to further processing or downstream computing were performed on alternative computing platforms, including Yunqi Academy of Engineering (Hangzhou, China) and Zhejiang Lab (Hangzhou, China).

## Competing Interests

1. Y. H., Z.-R. L., P. F., and J.-P. Y. have filed an application for a patent covering the work presented. The other authors declare no competing interests.

## Extended Data Figures

**Extended Data Figure 1:**
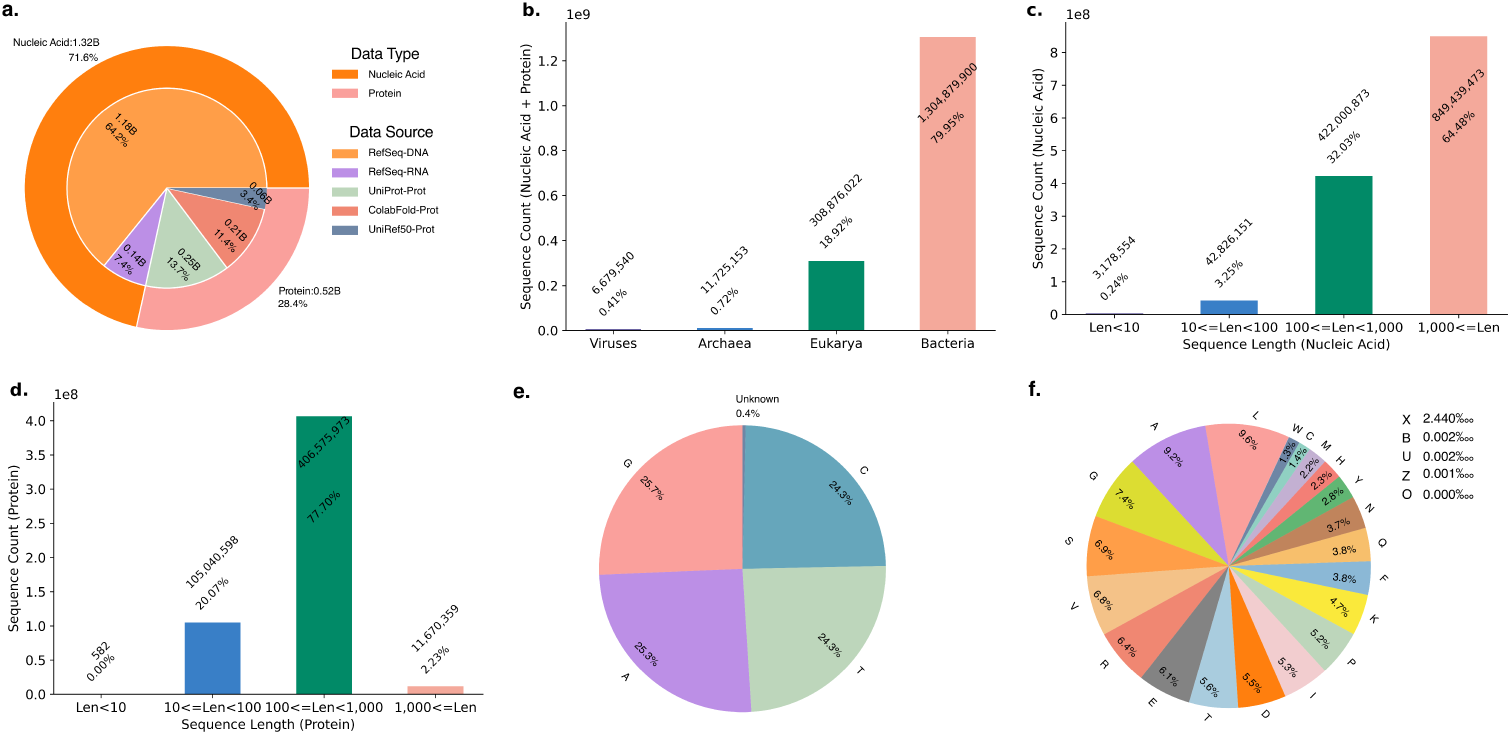
Overall statistics on pre-training data of LucaOne. **a.** Sequences (DNA, RNA, and proteins) were derived from RefSeq, UniProt, ColabFoldDB, and UniRef50. **b.** The data (nucleic acids and proteins) involved four superkingdom types: Viruses, Archaea, Eukarya, and Bacteria, of which Bacteria accounted for the most. **c.** The sequence length distribution of nucleic acids, with the most being more than 1,000. **d.** The sequence length distribution of proteins, with the maximum length ratio between 100 and 1,000. **e.** The proportion of five nucleotides (’A’, ’T’, ’C’, ’G’, and ’Unknown’) in nucleic acid sequences (’U’ compiled with ’T’ in RNA) and the four identified nucleotides were close in proportion. **f.** The proportion of the 20 standard amino acid letters and five other letters (including four non-standard amino acids and ’X’ for unknown amino acid) in the protein sequence, and Leucine has the highest proportion.

**Extended Data Figure 2:**
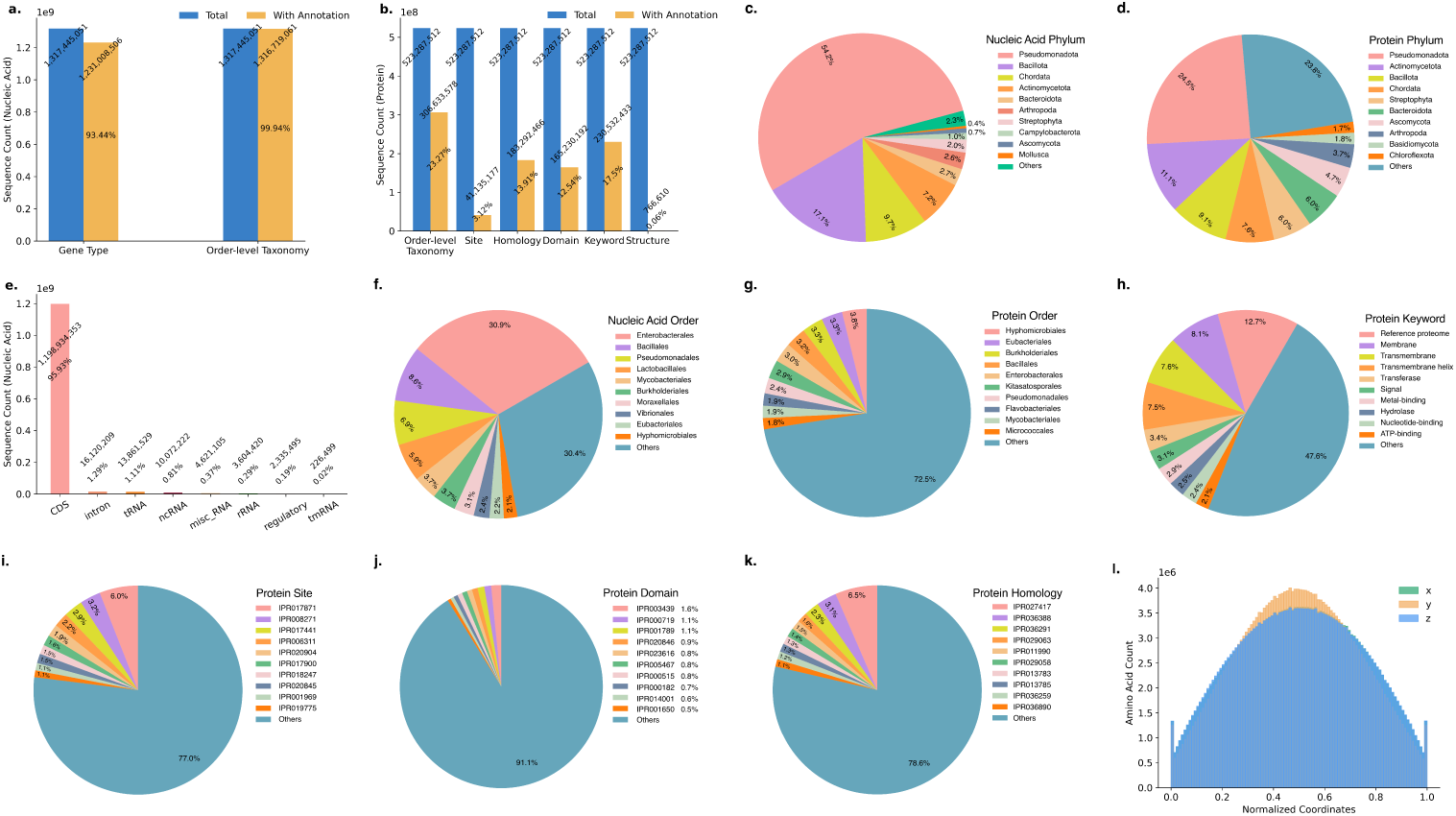
Annotation statistics on pre-training data of LucaOne. **a.** The proportion of genome region types and order-level taxonomy in nucleic acid. Most sequences have both types of annotation information. **b.** The proportion of the count of sequences with each of the selected six annotations, including order-level taxonomy, keyword, site, domain, homology, and tertiary structure, of which the proportion of sequence count with tertiary structure is tiny. **c. and d.** The proportion of sequence counts in the top 10 phylum-level taxonomy of nucleic acids and proteins, respectively. **e.** The distribution of eight selected genome region types in nucleic acids, of which the CDS region is the most. **f. and g.** The proportion of sequence counts in the top 10 order-level taxonomy (total 2,196 categories) of nucleic acids and proteins, respectively. **h.**∼**k.** The proportion of protein sequence counts in the top 10 keywords (total 1,179 categories), the top 10 site types (total 946 categories), the top 10 domain types (total 13,717 categories), and the top 10 homology types (total 3,442 categories), respectively. **l.** The *coord-(x, y, z)* distribution of *C_α_-atom* position (local normalization within a protein chain). It is very similar to the normal distribution. The distribution has a long tail in **c.**∼**f.**. The distribution is ladder decreasing in **g.**∼**k.**.

**Extended Data Figure 3:**
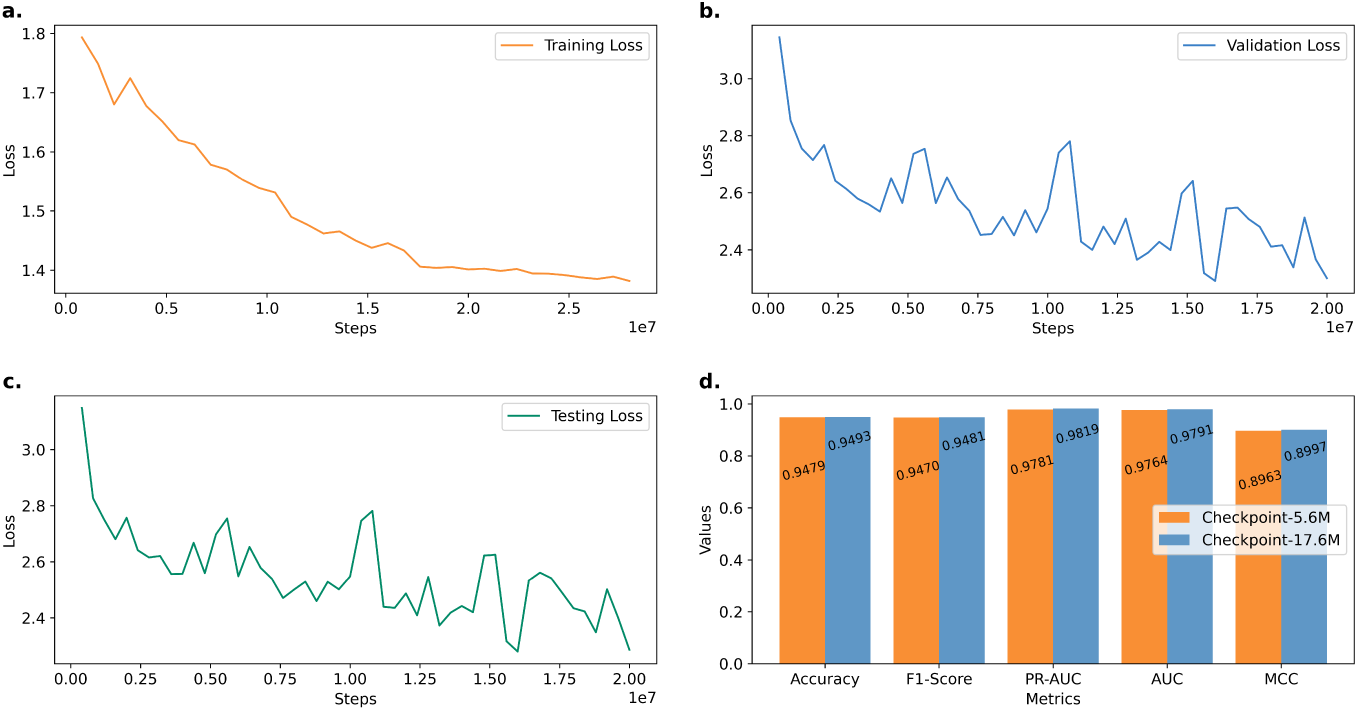
LucaOne 17.6M Checkpoint Selection Criteria. **a.** The loss trend during training. **b. and c.** The trend of loss on the validation and testing sets. **d.** Performance evaluation between 5.6M and 17.6M checkpoints on downstream tasks - ncRPI.

**Extended Data Figure 4:**
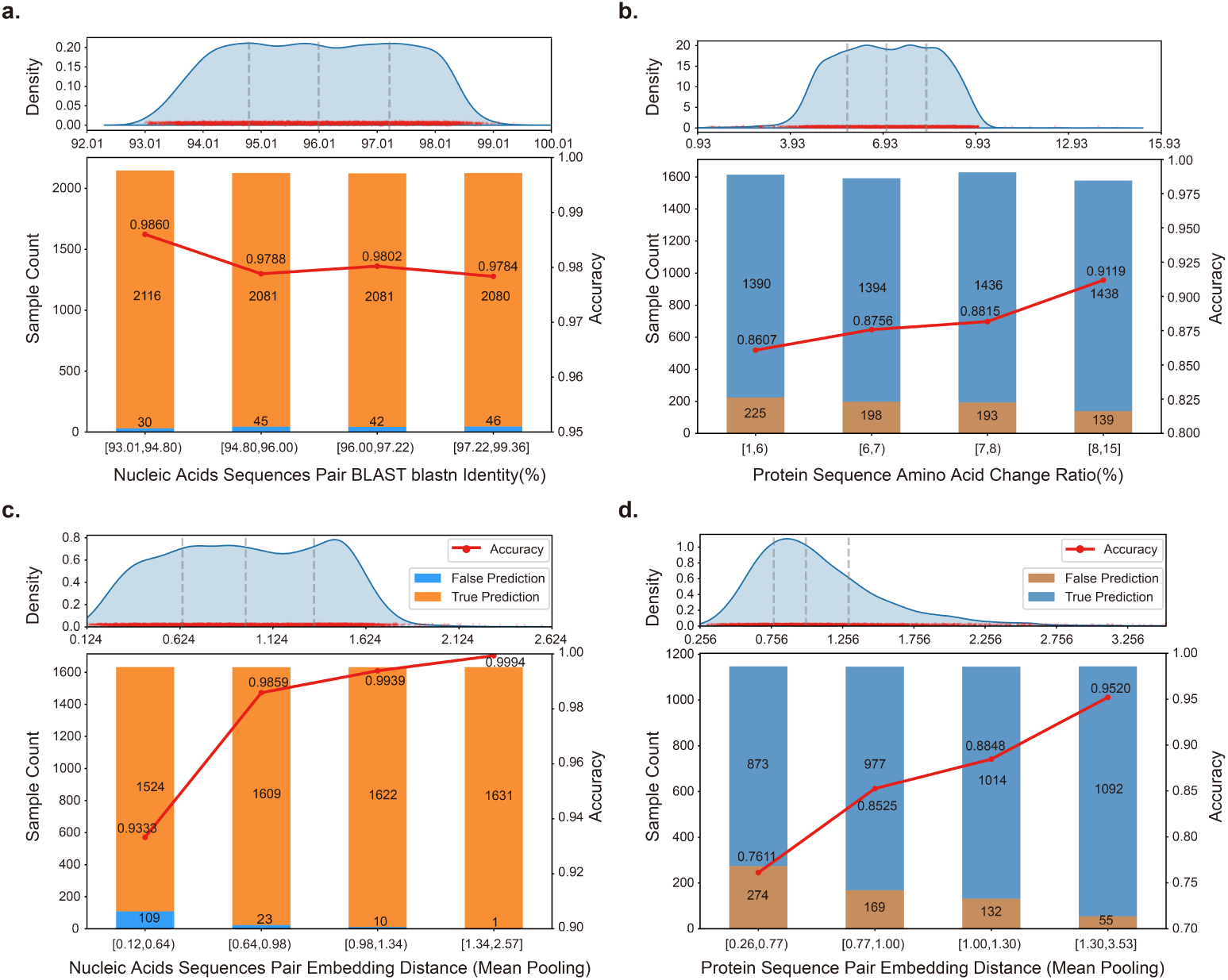
Identity Between Positive and Negative Samples and Prediction Accuracy in the Central Dogma Task. a. and. **b.** The relationship between sequence identity metrics and LucaOne model prediction accuracy: NCBI blastn sequence identity for nucleic acid and protein sequences before and after mutation. **c. and d.** Embedding Euclidean distances based on mean pooling and their prediction accuracy in LucaOne for nucleic acid and protein sequences before and after mutation. **Upper panels:** Sample distributions across sequence similarity, change ratio, or embedding Euclidean distance ranges. **Lower panels:** Prediction counts and accuracy of the LucaOne embedding within each respective range. **Note:** Data for **a. and b.** includes all nucleic acid and protein-negative samples from the validation and testing sets. Data for **c. and d.** includes only positive-negative sample pairs that are both present in the combined validation and testing datasets. Divide the statistical intervals of the metrics into quartiles according to the data distribution.

**Extended Data Figure 5:**
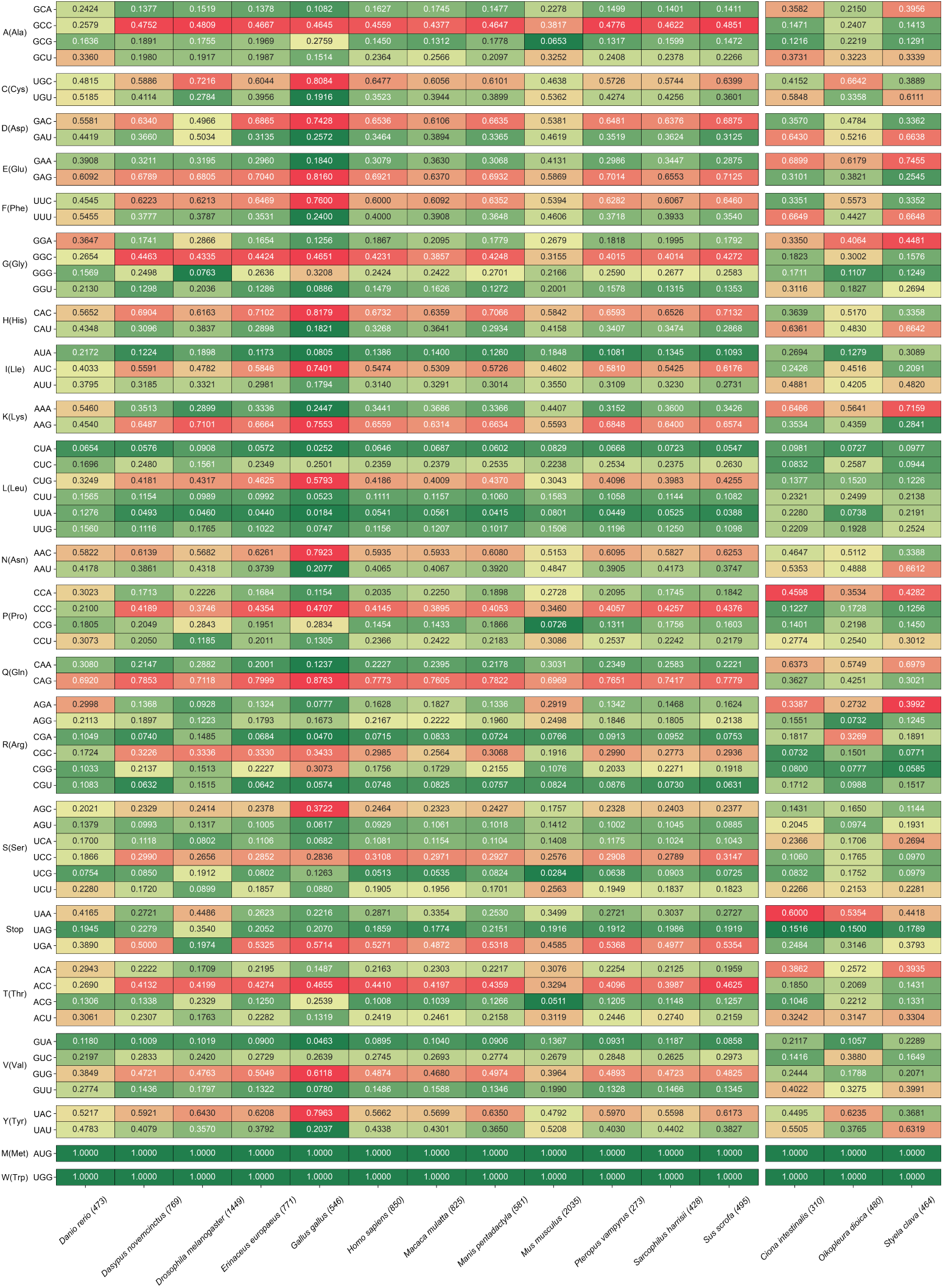
Codon Usage Heatmap. Based on the dataset for 15 species - Original Dataset plus two urochordates species. The distribution of different codons for a single amino acid totals 100%. Coloured representations indicate the relative proportions, where red signifies higher proportions and green signifies lower proportions.

## Extended Data Tables

**Extended Data Table 1:** Clustering metrics. The clustering scores (using K-Means++) of the four embedding methods on the S1, S2, and S3 datasets.

| DataSet | Embedding | ARI | AMI | HS | CS | V-measure | FMI |
| --- | --- | --- | --- | --- | --- | --- | --- |
| 12-Marine Species (S1) | Multi-OneHot | 0.1258 | 0.2782 | 0.2918 | 0.2949 | 0.2934 | 0.1998 |
|  | DNABert2 | 0.1313 | 0.2556 | 0.2686 | 0.2743 | 0.2714 | 0.2063 |
|  | LucaOne | **0.3410** | **0.5524** | **0.5488** | **0.5760** | **0.5621** | **0.4040** |
| 12-Clan Pfam (S2) | Multi-OneHot | 0.1062 | 0.1860 | 0.1763 | 0.2167 | 0.1944 | 0.2341 |
|  | ESM2-3B | 0.0731 | 0.1526 | 0.1617 | 0.1595 | 0.1606 | 0.1644 |
|  | LucaOne | **0.2379** | **0.4179** | **0.4228** | **0.4240** | **0.4234** | **0.3163** |
| 12-Go Terms (S3) | Multi-OneHot | 0.0942 | 0.1920 | 0.1934 | 0.2315 | 0.2107 | 0.2004 |
|  | ESM2-3B | 0.1079 | 0.2582 | 0.2596 | 0.2928 | 0.2752 | 0.2027 |
|  | LucaOne | **0.1535** | **0.3779** | **0.3707** | **0.4159** | **0.3920** | **0.2474** |

\* Bold values indicate the best performance for each dataset in the respective column.

**Extended Data Table 2:** Performance comparison for LucaOne and other embedding tools. LucaOne was not only compared with several existing embedding methods but also with itself, which was trained using nucleic acids and proteins separately (LucaOne-Gene/LucaOne-Prot). LucaOne, with mixing training, obtained the best performance for both the original dataset and the CDS-Protein dataset.

| Embedding | Network | Accuracy |  | F1-Score |  | AUC |  |
| --- | --- | --- | --- | --- | --- | --- | --- |
|  |  | Original | CDS-Protein | Original | CDS-Protein | Original | CDS-Protein |
| One-Hot | Transformer<br>+ Pooling<br>+ FC | 0.6667 | - | 0.0000 | - | 0.4943 | - |
| Random Init | Transformer<br>+ Pooling<br>+ FC | 0.6662 | - | 0.0015 | - | 0.5085 | - |
| DNABert2<br>+ ESM2-3B | Pooling<br>+ FC | 0.7309 | 0.7400 | 0.5689 | 0.6661 | 0.7491 | 0.7959 |
| LucaOne<br>(Gene/Prot) | Pooling<br>+ FC | 0.8048 | 0.8163 | 0.6804 | 0.7806 | 0.8616 | 0.9055 |
| LucaOne | Pooling<br>+ FC | **0.8453** | **0.8451** | **0.7392** | **0.8089** | **0.9101** | **0.9298** |

\* Bold values indicate the best score in the respective metric.

**Extended Data Table 3:**
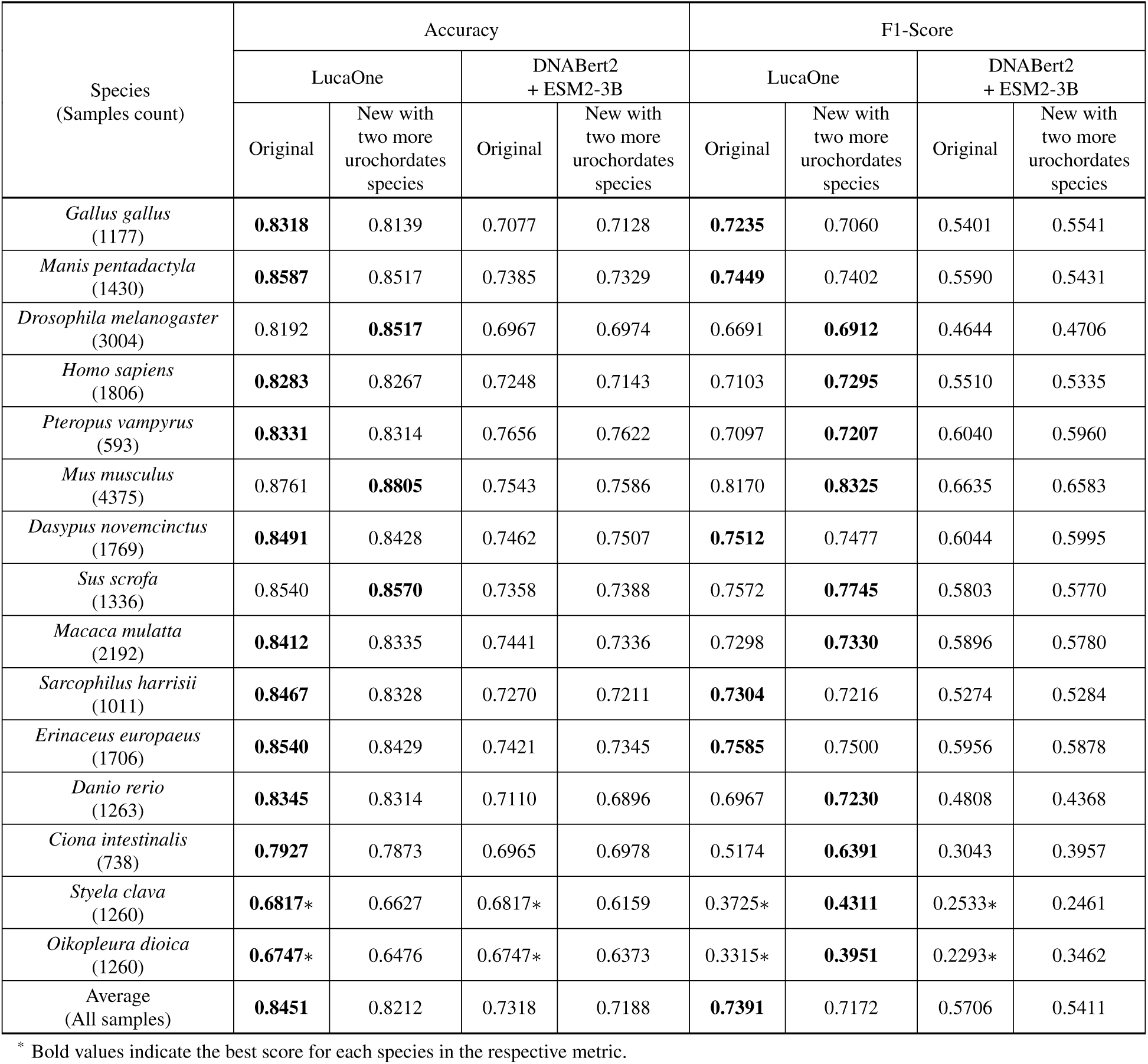
Comparative Performance Analysis. Comparative performance analysis (validation and testing set) of the models across diverse species datasets (Sample counts in brackets). The original dataset includes 13 species, and the new dataset added two more urochordates species data. F1-score and accuracy are calculated and presented. The top right ∗ indicates the predictive performances of the model trained by the original version of the Central Dogma dataset (w/o *Oikopleura Dioica* and *Styela Clava* data). More details in **Data Availability**.

| Species<br>(Samples count) | Accuracy |  |  |  | F1-Score |  |  |  |
| --- | --- | --- | --- | --- | --- | --- | --- | --- |
|  | LucaOne |  | DNABert2<br>+ ESM2-3B |  | LucaOne |  | DNABert2<br>+ ESM2-3B |  |
|  | Original | New with<br>two more<br>urochordates<br>species | Original | New with<br>two more<br>urochordates<br>species | Original | New with<br>two more<br>urochordates<br>species | Original | New with<br>two more<br>urochordates<br>species |
| *Gallus gallus*<br>(1177) | **0.8318** | 0.8139 | 0.7077 | 0.7128 | **0.7235** | 0.7060 | 0.5401 | 0.5541 |
| *Manis pentadactyla*<br>(1430) | **0.8587** | 0.8517 | 0.7385 | 0.7329 | **0.7449** | 0.7402 | 0.5590 | 0.5431 |
| *Drosophila melanogaster*<br>(3004) | 0.8192 | **0.8517** | 0.6967 | 0.6974 | 0.6691 | **0.6912** | 0.4644 | 0.4706 |
| *Homo sapiens*<br>(1806) | **0.8283** | 0.8267 | 0.7248 | 0.7143 | 0.7103 | **0.7295** | 0.5510 | 0.5335 |
| *Pteropus vampyrus*<br>(593) | **0.8331** | 0.8314 | 0.7656 | 0.7622 | 0.7097 | **0.7207** | 0.6040 | 0.5960 |
| *Mus musculus*<br>(4375) | 0.8761 | **0.8805** | 0.7543 | 0.7586 | 0.8170 | **0.8325** | 0.6635 | 0.6583 |
| *Dasypus novemcinctus*<br>(1769) | **0.8491** | 0.8428 | 0.7462 | 0.7507 | **0.7512** | 0.7477 | 0.6044 | 0.5995 |
| *Sus scrofa*<br>(1336) | 0.8540 | **0.8570** | 0.7358 | 0.7388 | 0.7572 | **0.7745** | 0.5803 | 0.5770 |
| *Macaca mulatta*<br>(2192) | **0.8412** | 0.8335 | 0.7441 | 0.7336 | 0.7298 | **0.7330** | 0.5896 | 0.5780 |
| *Sarcophilus harrisii*<br>(1011) | **0.8467** | 0.8328 | 0.7270 | 0.7211 | **0.7304** | 0.7216 | 0.5274 | 0.5284 |
| *Erinaceus europaeus*<br>(1706) | **0.8540** | 0.8429 | 0.7421 | 0.7345 | **0.7585** | 0.7500 | 0.5956 | 0.5878 |
| *Danio rerio*<br>(1263) | **0.8345** | 0.8314 | 0.7110 | 0.6896 | 0.6967 | **0.7230** | 0.4808 | 0.4368 |
| *Ciona intestinalis*<br>(738) | **0.7927** | 0.7873 | 0.6965 | 0.6978 | 0.5174 | **0.6391** | 0.3043 | 0.3957 |
| *Styela clava*<br>(1260) | **0.6817\*** | 0.6627 | 0.6817* | 0.6159 | 0.3725* | **0.4311** | 0.2533* | 0.2461 |
| *Oikopleura dioica*<br>(1260) | **0.6747\*** | 0.6476 | 0.6747* | 0.6373 | 0.3315* | **0.3951** | 0.2293* | 0.3462 |
| Average<br>(All samples) | **0.8451** | 0.8212 | 0.7318 | 0.7188 | **0.7391** | 0.7172 | 0.5706 | 0.5411 |

\* Bold values indicate the best score for each species in the respective metric.

**Extended Data Table 4:**
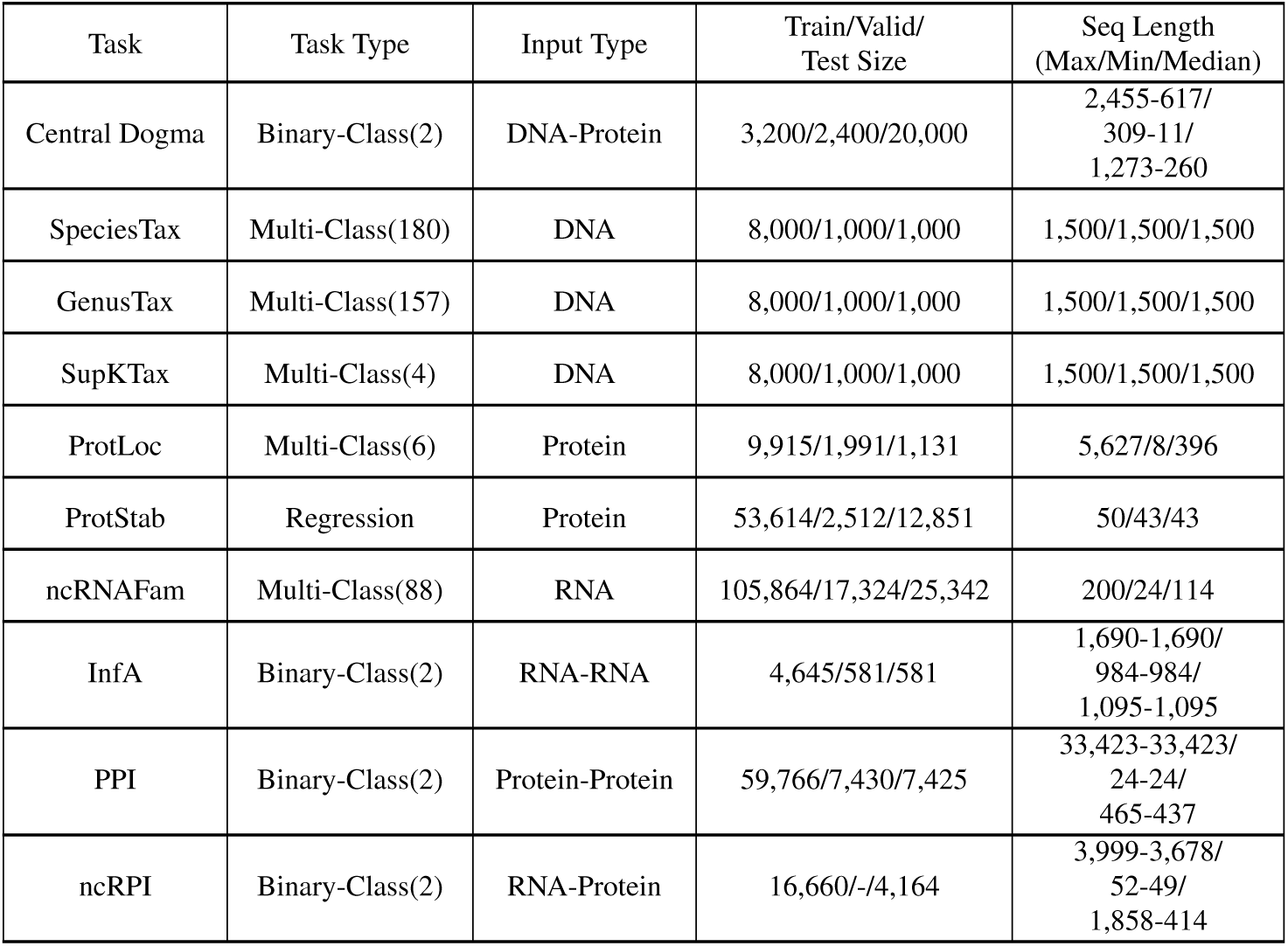
Details on downstream validation tasks. Details of 10 downstream tasks, including task name, task type, input type of task, sample number of training set, validation set, test set, and sequence length statistics of each task.

**Extended Data Table 5:**
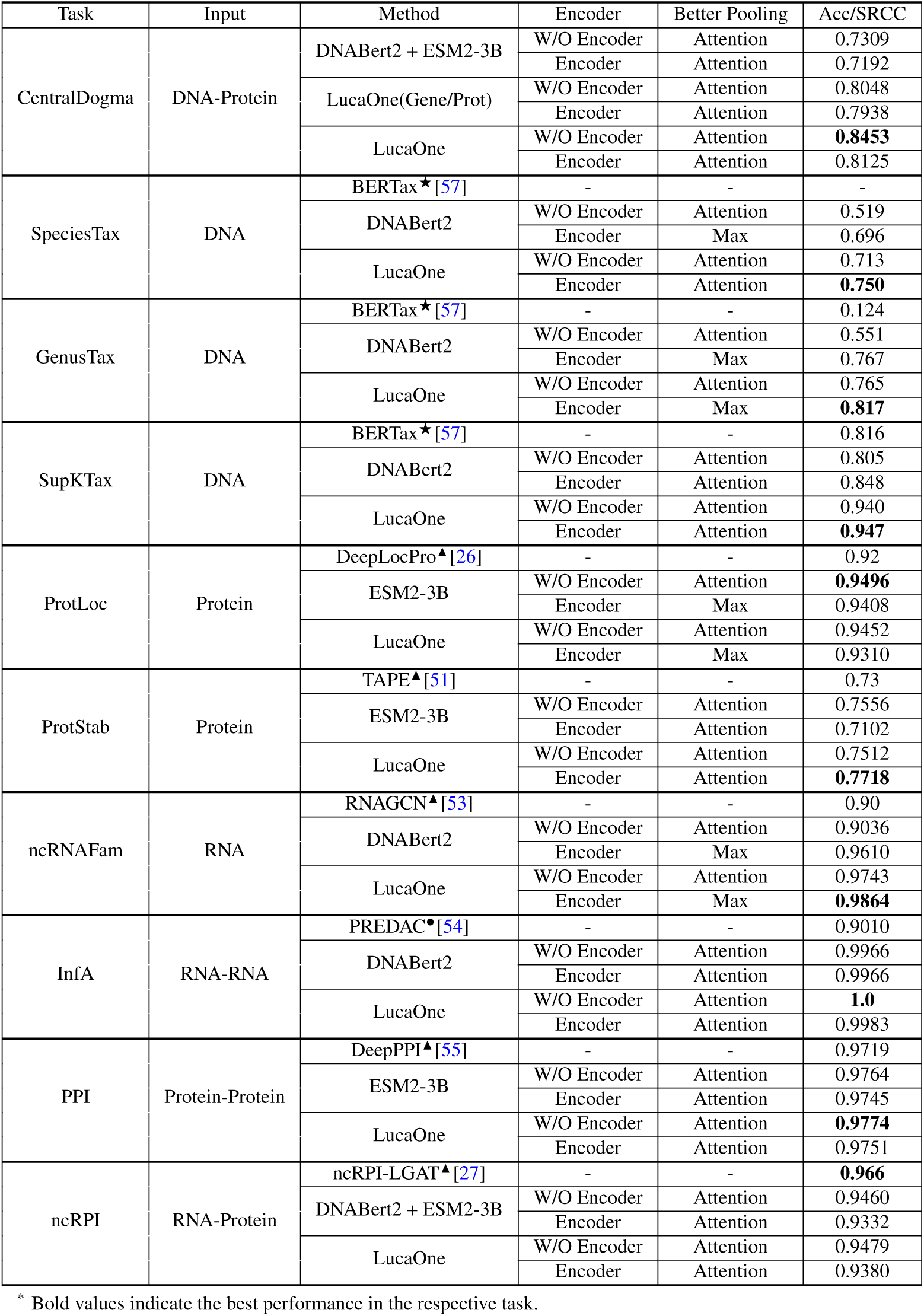
Detailed results of the testing set on downstream validation tasks (results of the better pooling method for each task with or without encoder). The top right ★ indicates inference using the trained method, the top right ▴ indicates direct use of the results in its paper, and the top right • indicates repetition using its method and higher than the results in the paper.

## Notes

http://47.93.21.181/lucaone/

https://github.com/LucaOne/LucaOne

https://doi.org/10.5281/zenodo.15171943

### Competing Interest Statement

Yong He, Zhaorong Li, Pan Fang, and Jieping Ye have filed an application for a patent covering the work presented. The other authors declare no competing interests.

### Summary of Updates

Additional details on model training, comparative experiments, and other supplementary information have been provided in the Supplementary Notes. Figure 3 has been significantly updated.

