## Supplementary Note for "Generalized Biological Foundation Model with Unified Nucleic Acid and Protein Language"

### Supplementary Notes

**2 GO-guided Functional Characterization Through Embedding Analysis** **4**

**3 Analysis of Misclassified Samples on Central Dogma Task** **6**

**4 Data Details for CDS-Protein Task** **7**

**5 Task for Cross-Species Homologous Gene Pairs** **7**

**6 Convergence of Nucleic Acid and Protein Sequences for the Same Gene** **8**

**7 Task on Pseudogene Correction** **9**

**8 Task on ProtLoc with Nucleic Acid Embeddings** **11**

**9 Task on SYN and NON-SYN Mutations Embeddings Distances** **11**

**10 Task on Nearest Neighbors Analysis for Urochordates Species** **12**

### 1 Loss Functions for Pretraining Tasks

#### 1.1 Token Level Tasks

The loss functions $\mathcal{L}_{GeneMask}$ and $\mathcal{L}_{ProtMask}$ of the two token-level mask language pretraining tasks are defined as follows:

$$\mathcal{L}_{GeneMask} = -\frac{1}{|s_1|}\sum_{i\in s_1} log[\hat{p}^1(nt_i|nt\backslash i)] \tag{1}$$

$$\mathcal{L}_{ProtMask} = -\frac{1}{|s_2|}\sum_{i\in s_2} log[\hat{p}^2(aa_i|aa\backslash i)] \tag{2}$$

where $s_1$ and $s_2$ are the sets of masked nucleotides and amino acids, respectively, and $\hat{p}^1(nt_i|nt\backslash i)$ and $\hat{p}^2(aa_i|aa\backslash i)$ indicate the probability of the masked nucleotide $nt_i$ and the masked amino acid $aa_i$ based on their respective contexts $(nt\backslash i)$ and $(aa\backslash i)$, respectively.

#### 1.2 Span Level Tasks

The loss functions $\mathcal{L}_{GenomeType}$, $\mathcal{L}_{ProtSite}$, $\mathcal{L}_{ProtHomo}$ and $\mathcal{L}_{ProtDomain}$ of the four span-level pre-training tasks are as follows:

$$\mathcal{L}_{GenomeType} = -\frac{1}{|s_3|}\sum_{i\in s_3} log[\hat{p}^3(nt_i|nt\backslash i)] \tag{3}$$

$$\mathcal{L}_{ProtSite} = -\frac{1}{|s_4|}\sum_{i\in s_4} log[\hat{p}^4(aa_i|aa\backslash i)] \tag{4}$$

$$\mathcal{L}_{ProtHomo} = -\frac{1}{|s_5|}\sum_{i\in s_5} log[\hat{p}^5(aa_i|aa\backslash i)] \tag{5}$$

$$\mathcal{L}_{ProtDomain} = -\frac{1}{|s_6|}\sum_{i\in s_6} log[\hat{p}^6(aa_i|aa\backslash i)] \tag{6}$$

where $s_3$, $s_4$, $s_5$, and $s_6$ represent nucleotide positions set in which eight selected genome type labels exist, amino acid positions set in which 946 selected protein sites exist, amino acid positions set in which 3,442 selected protein homology categories exist, and amino acid positions set in which 13,717 selected protein domain classes exist, respectively. $\hat{p}^3(nt_i|nt\backslash i)$indicates the probability of the category of the nucleotide genome region $nt_i$, and $\hat{p}^4(aa_i|aa\backslash i)$, $\hat{p}^5(aa_i|aa\backslash i)$and $\hat{p}^6(aa_i|aa\backslash i)$ indicate the probability of the category of the site, the homologous category and the domain category of the amino acid $aa_i$ based on its context $(aa\backslash i)$, respectively.

#### 1.3 Seq Level Tasks

The loss functions $\mathcal{L}_{GeneTaxonomy}$, $\mathcal{L}_{ProtTaxonomy}$, and $\mathcal{L}_{ProtKeyword}$ of the three sequence-level pre-training tasks are as follows:

$$\mathcal{L}_{GeneTaxonomy} = -\frac{1}{|s_7|}\sum_{i\in s_7} log[\hat{p}^7(seq_i)] \tag{7}$$

$$\mathcal{L}_{ProtTaxonomy} = -\frac{1}{|s_8|}\sum_{i\in s_8} log[\hat{p}^8(seq_i)] \tag{8}$$

$$\mathcal{L}_{ProtKeyword} = -\frac{1}{|s_9|}\sum_{i\in s_9}\frac{1}{|kws_i|}\sum_{j\in kws_i} log[\hat{p_j}^9(seq_i)] \tag{9}$$

where $s_7$, $s_8$, and $s_9$ represent the set of nucleic acid sequences in which the order-level taxonomy label exists, the set of protein sequences in which the order-level taxonomy label exists, and the set of protein sequences in which the keyword exists, respectively. $kws_i$ is the set of keywords in the protein sequence $i$. $\hat{p}^7(seq_i)$ indicates the probability of the category of order-level taxonomy of the nucleic acid sequence $i$. $\hat{p}^8(seq_i)$ and $\hat{p_j}^9(seq_i)$ indicate the probability of the order-level taxonomy category and the keyword $j$ of the protein sequence $i$, respectively.

#### 1.4 Structure Level Tasks

The loss function $\mathcal{L}_{ProtStructure}$ of the amino acid position regression structure-level pre-training task is as follows:

$$\mathcal{L}_{ProtStructure} = \frac{1}{3*|s_{10}|}\sum_{i\in s_{10}}\sum_{p^{10}\in\{x,y,z\}} |p^{10}(aa_i) - \hat{p}^{10}(aa_i|aa\backslash i)| \tag{10}$$

where $s_{10}$ is the set of amino acid positions in which the *coord-(x,y,z)* exists, $p^{10}(aa_i)$ is the local normalized *coord-(x,y,z)* of the $C_\alpha$*-atom* in the amino acid, and $\hat{p}^{10}(aa_i|aa\backslash i)$ indicates the prediction *coord-(x,y,z)* of the amino acid $aa_i$ based on its context $(aa\backslash i)$.

#### 1.5 Weighted Combined Loss

We employed 10 different pretraining tasks, assigning an equal weight of 1.0 to Gene-Mask, Prot-Mask, Prot-Keyword, and Prot-Structure tasks, while assigning a reduced weight of 0.2 to the remaining tasks to equilibrate task complexity with the range of their loss function values.

$$\mathcal{L} = 1.0\mathcal{L}_{GeneMask} + 1.0\mathcal{L}_{ProtMask} + 0.2\mathcal{L}_{GenomeType} + 0.2\mathcal{L}_{GeneTaxonomy}$$

$$+0.2\mathcal{L}_{ProtSite} + 0.2\mathcal{L}_{ProtHomo} + 0.2\mathcal{L}_{ProtDomain} + 0.2\mathcal{L}_{ProtTaxonomy}$$
$$+1.0\mathcal{L}_{ProtKeyword} + 1.0\mathcal{L}_{ProtStructure} \quad (11)$$

### 2 GO-guided Functional Characterization Through Embedding Analysis

To further evaluate the efficacy of LucaOne embeddings in Gene Ontology (GO) annotation tasks, we constructed two comprehensive datasets, each containing three distinct subsets corresponding to biological processes (BP), cellular components (CC), and molecular functions (MF).

**S3-GO-V1 Dataset Construction**: Adhering to the methodology outlined in the main text, we identified the top 12 GO terms within each category (BP, CC, MF) based on the following criteria: **(a)** exclusion of ancestor terms, **(b)** minimum GO hierarchy level of 4, and **(c)** representation among the top 12 most frequent terms in the original dataset. From each selected GO term, we randomly sampled 100 sequences with lengths ranging from 100 to 2,500 amino acids, ensuring the exclusion of sequences spanning multiple GO terms. This process yielded three distinct subsets: S3-GO-V1-BP (also utilized in the main text), S3-GO-V1-CC, and S3-GO-V1-MF. We subsequently applied MultiHot encoding, ESM2-3B embeddings, and LucaOne embeddings to these subsets, followed by t-SNE visualization and KMeans++ clustering analysis. The clustering performance was evaluated using six established metrics (**Supplementary Figure 5 and 7**). Notably, LucaOne embeddings demonstrated superior performance across all six clustering metrics in all three datasets, with particularly significant advantages observed in the S3-GO-V1-BP and S3-GO-V1-CC subsets.

**S3-GO-V2 Dataset Construction**: To address potential limitations in discriminative capability within S3-GO-V1, which may arise from inherent interrelationships among GO terms, we developed an enhanced dataset version specifically designed to improve differentiation in t-SNE visualization and clustering analyses. For each GO category (BP, CC, MF), we implemented a rigorous selection process to identify 12 GO terms with minimal semantic correlation, designated as S3-GO-V2. The selection criteria included: **(a)** exclusion of ancestor terms, **(b)** minimum GO hierarchy level of 4, and **(c)** availability of more than 100 associated sequences. We employed a greedy algorithm to maximize term dissimilarity based on Lin similarity. Specifically, the selection process was initiated with a randomly chosen seed GO term, followed by iterative selection of the term exhibiting the lowest Lin similarity to the current set of selected terms, continuing until 12 terms were identified.

For each selected GO term, we randomly sampled 100 sequences with lengths between 100 and 2,500 amino acids, ensuring the exclusion of sequences annotated with multiple GO terms. This procedure generated three distinct subsets: S3-GO-V2-BP, S3-GO-V2-CC, and S3-GO-V2-MF. We subsequently applied MultiHot encoding, ESM2-3B embeddings, and LucaOne embeddings to these subsets, followed by t-SNE visualization and KMeans++ clustering analysis. The clustering performance was evaluated using six established metrics (**Supplementary Figure 6 and 7**). LucaOne embeddings demonstrated optimal performance across all six clustering metrics.

**Correlation Analysis Between GO Term Similarity and Sequence Embedding Similarity**: Furthermore, we conducted an additional investigation to validate whether embedding similarity between protein sequences across different GO annotations correlates with the inherent dissimilarities of their respective GO terms. Specifically, we examined whether sequence clusters with greater functional divergence demonstrate proportionally larger distances in their embedding representations. Employing the identical methodology as described for S3-GO-V1, we systematically selected the top 12 Gene Ontology (GO) terms within the cellular component category, yielding a comprehensive dataset of 1,200 sequences across these 12 GO terms. Direct inter-term correlations were quantified using Lin similarity (**Eq. (12)**), generating a correlation matrix $M_1^{12\times 12}$, where each element $m_{i,j} \in [0,1]$. To investigate sequence embedding-based correlations, we implemented three distinct computational approaches:

**(a)** Construction of a cosine similarity matrix derived from the centroids of sequence embeddings for each respective GO term (**Eq. (13)**);

**(b)** Computation of mean pairwise cosine similarity values across sequence embeddings within each GO term pair (**Eq. (14)**);

**(c)** Determination of median pairwise cosine similarity values across sequence embeddings within each GO term pair (**Eq. (15)**).

These methodologies yielded three corresponding matrices: $M_{21}^{12\times 12}$, $M_{22}^{12\times 12}$, and $M_{23}^{12\times 12}$, with cosine similarity values normalized into $[0,1]$ through the transformation $\frac{1+cosine}{2}$. Subsequently, we calculated both the Pearson and Spearman correlation coefficients, accompanied by their respective p-values, between $M_1^{12\times 12}$ and each of the embedding-derived matrices $M_{21}^{12\times 12}$, $M_{22}^{12\times 12}$, and $M_{23}^{12\times 12}$ (**Supplementary Figure 8**). The analytical results demonstrate significant coherence between direct GO term correlations and those derived from sequence embedding analyses. Notably, LucaOne embeddings exhibited superior correlation in comparison to ESM2-3B embeddings across all evaluated metrics.

$$
\begin{gathered}
M_1^{12\times 12} = M_{LinSim} = \begin{pmatrix} m_1^1 & \cdots & m_1^{12} \\ \vdots & \vdots & \vdots \\ m_{12}^1 & \cdots & m_{12}^{12} \end{pmatrix} \\
m_i^j = LinSim(GO_i, GO_j) = \frac{2 \times IC(LCS(GO_i, GO_j))}{IC(GO_i) + IC(GO_j)} \\
m_i^j \in [0,1]
\end{gathered} \tag{12}
$$

where $IC(GO_i)$ represents the information content of $GO_i$, which is usually calculated as the negative logarithm of the probability of encountering $GO_i$, i.e., $IC(GO_i) = -log(p(GO_i))$, and $LCS(GO_i, GO_j)$ represents the Least Common Subsumer of $GO_i$ and $GO_j$. $LCS$ is the most specific ancestor node that is common to both $GO_i$ and $GO_j$ in the ontology hierarchy.

$$
\begin{gathered}
M_{21}^{12\times 12} = M_{CentroidsCosine} = \begin{pmatrix} m_1^1 & \cdots & m_1^{12} \\ \vdots & \vdots & \vdots \\ m_{12}^1 & \cdots & m_{12}^{12} \end{pmatrix} \\
m_i^j = (1 + Cosine(GO_i, GO_j))/2 = (1 + Cosine(\frac{\sum_{p=1}^{100} e_{ip}}{100}, \frac{\sum_{q=1}^{100} e_{jq}}{100}))/2 \\
p, q \in \{1, 2, ..., 100\} \\
m_i^j \in [0, 1]
\end{gathered}
\tag{13}
$$

$$
\begin{gathered}
M_{22}^{12\times 12} = M_{MeanPairwiseCosine} = \begin{pmatrix} m_1^1 & \cdots & m_1^{12} \\ \vdots & \vdots & \vdots \\ m_{12}^1 & \cdots & m_{12}^{12} \end{pmatrix} \\
m_i^j = (1 + Cosine_{mean}(GO_i, GO_j))/2 = (1 + \frac{\sum_{p=1}^{100} \sum_{q=1}^{100} Cosine(e_{ip}, e_{jq})}{100 \times 100})/2 \\
p, q \in \{1, 2, ..., 100\} \\
m_i^j \in [0, 1]
\end{gathered}
\tag{14}
$$

$$
\begin{gathered}
M_{23}^{12\times 12} = M_{MedianPairwiseCosine} = \begin{pmatrix} m_1^1 & \cdots & m_1^{12} \\ \vdots & \vdots & \vdots \\ m_{12}^1 & \cdots & m_{12}^{12} \end{pmatrix} \\
m_i^j = (1 + Cosine_{median}(GO_i, GO_j))/2 = (1 + Median\{Cosine(e_{ip}, e_{jq})\})/2 \\
p, q \in \{1, 2, ..., 100\} \\
m_i^j \in [0, 1]
\end{gathered}
\tag{15}
$$

where $GO_i$ represents the $i_{th}$ GO term, $e_{ip}$ represents the embedding vector of the $p_{th}$ sequence sampled under $GO_i$ (the embedding matrix of the sequence is transformed into an embedding vector by mean pooling strategy), and $Cosine$ represents the cosine similarity between two vectors. Each GO term randomly samples 100 sequences.

All datasets for this analysis are available; please see Section - **Data Availability-Supplementary**.

### 3 Analysis of Misclassified Samples on Central Dogma Task

We constructed two negative samples for each positive sample by inserting, replacing, or deleting nucleotides or amino acids. Negative nucleic acid samples were generated through simulated substitution, insertion, and deletion operations. Mutations were restricted to gene sequences, excluding regions within 100 base pairs upstream and downstream. For each sequence, the number of mutation sites was determined by assigning a random mutation rate between 2% and 8%. Negative protein samples were generated by systematically altering amino acid residues at specific sites while preserving the overall sequence length. For each protein sequence, the percentage of residues subjected to alteration was randomly determined, generally ranging from 3% to 10%. Unique alteration sites were selected from the sequence at random, with the exception of the first residue, which remained unchanged to maintain consistency with natural protein initiation regions. **Extended Data Figure 4** shows details.

The comparison analysis of performance across different subsets of exon count was presented in **Supplementary Figure 9**.

### 4 Data Details for CDS-Protein Task

CDS sequences replaced the extended nucleic acid sequences in the original positive samples, forming a new set of positive samples while retaining only the CDS-Protein pairs that adhered to the triplet codon translation logic. For protein-altered negative samples, protein sequences were preserved, and their corresponding nucleic acid sequences were replaced with CDS sequences. Nucleotide-mutated negative samples were regenerated following the original strategy.

The training, validation, and testing set allocations from the original dataset were maintained. The final data set of CDS proteins paired comprises 24,422 samples, distributed as 2,422: 1,851: 15,304 in the training, validation, and testing sets (roughly 4: 3: 25). The other setting of the model was the same as in the original task. All datasets are available; please see Section - **Data Availability-Supplementary**.

### 5 Task for Cross-Species Homologous Gene Pairs

We designed an additional task related to the central dogma by modifying the negative samples in the original study. Instead of manually altering the sequences, the negative samples were replaced with homologous genes from closely related species. This approach is more challenging than randomly pairing nucleic acid and protein sequences, as the variations between homologous genes are more subtle.

**Data Details**: This cross-species dataset was constructed by pairing nucleic acid and protein sequences of the same gene across different species. Target genes were selected from mammalian genomes in the NCBI RefSeq genomes database at the representative and reference genome levels. These genes were identified based on their frequency of occurrence in mammalian species, with the top 50 genes that occur most frequently having nucleic acid sequences of less than 2,000 bp chosen as targets. Positive samples were generated by extracting nucleic acid sequences corresponding to each target gene, including 100 bp flanking regions upstream and downstream of the gene and their corresponding protein sequences. These pairs of nucleic acid and protein sequences formed the positive dataset. Negative samples were constructed by fixing the nucleic acid sequence of each positive pair and randomly pairing it with the protein sequence of the same gene from a different species. To control the size of the dataset while maintaining diversity, two negative samples were created for each positive sample, with mismatched protein sequences validated to confirm that they were distinct from the original protein sequence in the positive pair. The final positive dataset comprises 7,728 sequence pairs, spanning 50 genes and 170 species, with a positive-to-negative sample ratio of 1: 2. Following the same data allocation strategy as the original dataset, the positive and negative samples were randomly shuffled, divided into 32 subsets, and divided into training, validation, and testing sets in a 4: 3: 25 ratio.

Under the similar 4: 3: 25 data partition scheme, the model achieved a prediction accuracy of 55.71% on the testing set. When the dataset was repartitioned with an 8: 1: 1 ratio to increase the size of the training set, the model performance improved significantly, achieving an accuracy of 74.54%. LucaOne consistently outperformed the DNABert2 + ESM2-3B combination in both data allocation schemes. For date details and performance comparison, then see **Supplementary Figure 10** and Section - **Data Availability-Supplementary**.

### 6 Convergence of Nucleic Acid and Protein Sequences for the Same Gene

The following analysis demonstrated that, although nucleic acid and protein sequences were not paired during model training, nucleic acid and protein sequences corresponding to the same gene exhibited convergence in the LucaOne Embedding Space.

To better analyze this problem, a more comprehensive dataset with more genes was constructed for evaluation. Sampling was conducted from three kingdoms in the NCBI RefSeq genome database categorized as reference genomes: Bacteria, Eukaryota, and Viruses. Seven species were randomly selected from each category, and genes with nucleic acid sequence lengths shorter than 2,500 bp were selected. Both the nucleic acid sequences and corresponding protein sequences were aligned to the trained sequences of LucaOne checkpoint-5.6M using NCBI blastn and Diamond blastp, respectively. Any genes with alignment hits were excluded (resulting in 7,818 pairs), and only species retaining more than 10 genes after filtering were preserved (resulting in 7,804 pairs). For species containing more than 200 genes, 200 genes were randomly sampled; all sequences were retained for species with fewer than 200 genes. The final dataset comprised 1284 gene-protein pairs across 10 species (Dataset Statistics in **Supplementary Figure 11**).

Based on the embedding representations (utilizing the $[CLS]$ embedding vector) of 1,284 gene-protein pairs, we performed t-SNE dimensionality reduction for visualization and presentation (**Supplementary Figure 12-a**). The embedding representations of genes and proteins were effectively separated. At the gene level, distinct species were also clearly differentiated. For proteins, the inter-species discrimination was relatively weaker. Furthermore, the mean Euclidean distances between gene-protein pairs were consistently similar across different species.

**Analysis of the Jaccard Index:** The embedding utilized for Jaccard index computations is derived from the output of the final layer of the Transformer-Encoder architecture, specifically the embedding matrix with dimensions of the sequence length $\times$ 2560. This matrix undergoes the transformation into a vector representation through the application of a mean pooling strategy, which is subsequently employed for Euclidean distance computations.

In this dataset, each gene-protein pair is uniquely identified by a sample ID (unique_id, gene_seq, prot_seq). For every gene-protein pair $(id_i, gene\_seq_i, prot\_seq_i)$, we independently computed the Jaccard similarity, followed by the calculation of the mean Jaccard score across all pairs using **Eq. (16)**:

$$Mean\ Jaccard = \frac{1}{N}\sum_{i=1}^{N}\frac{|S_i^{Gene} \cap S_i^{Prot}|}{|S_i^{Gene} \cup S_i^{Prot}|} \in [0,1] \tag{16}$$

where, $N$ is the number of Gene-Protein pairs, $i$ is the index of pairs, $S_i^{Gene}$ is the top $K$ nearest neighbors of the gene in $Pair_i$, and $S_i^{Prot}$ is the top $K$ nearest neighbors of the protein in $Pair_i$.

We conducted a comprehensive comparison based on the Jaccard Index evaluation among LucaOne embedding approaches, global/local sequence alignment, and DNABert2 + ESM2-3B embedding. For the sequence alignment analysis, we implemented global and local alignment using the 'pairwise2.align.globalxx' and 'pairwise2.align.localxx' functions from the Biopython package (**Eq. (17)**).

$$Seq\ Align\ Sim = \frac{Score_{BestAlign}}{max\{|seq_a|, |seq_b|\}} \in [0,1] \tag{17}$$

where, $Score_{BestAlign}$ is the score of the best alignment of $Seq_a$ and $Seq_b$, and $|seq|$is the length of the sequence.

We subsequently calculated the mean Jaccard similarity for the top 10 matches (TOP 1$\sim$10) across these three methodologies (**Supplementary Figure 12-b**; note that the global and local alignment methods yielded identical results in this analysis).

The comparative analysis revealed that LucaOne consistently outperformed the alternative methods. Notably, while DNABert2 + ESM2-3B represents an integrated approach combining separate nucleic acid and protein models, its performance was inferior to traditional sequence alignment methods. This finding suggests that joint training of nucleic acid and protein models, as implemented in LucaOne, provides superior performance for integrated nucleic acid-protein analysis scenarios. All datasets are available; please see Section - **Data Availability-Supplementary**.

### 7 Task on Pseudogene Correction

We conducted a mask task prediction analysis (zero-shot) on true gene (protein-coding) and pseudogene pairs data. The analysis includes three specific experiments:

**(a) Pseudogene Correction:** The mismatched nucleotides are replaced by '[MASK]' for each pseudogene sequence. Then, LucaOne is applied to predict the correct nucleotides corresponding to the true gene. The accuracy of restoring the pseudogene to match the true gene nucleotides is calculated. In this scenario, the input is the pseudogene sequence with masked mismatched positions, and the ground truth is the corresponding true gene sequence.

**(b) Pseudogene Keep:** The mismatched nucleotides are replaced by '[MASK]' for each pseudogene sequence. LucaOne is then applied to predict the nucleotides, maintaining them as pseudogene nucleotides. The accuracy of preserving the pseudogene's nucleotides is measured. In this scenario, the input is the pseudogene sequence with masked mismatched positions, and the prediction target is the pseudogene sequence.

**(c) Gene Random Mask Recover:** A random sampling is applied to mask nucleotides for each true gene sequence. The proportion of masked nucleotides matches the mismatch rate observed in the corresponding pseudogene. LucaOne is then applied to predict and restore the masked true gene nucleotides. The input is a true gene sequence with randomly masked positions, and the ground truth is the true gene sequence.

For each masked position, nucleotide prediction involves a multi-class classification from a vocabulary of 39 tokens. For this task, our hypothesis is that if LucaOne able to distinguish between real genes and pseudogenes, the Gene Random Mask Recover Rate should exceed the Pseudogene Keep Rate, as the former's input comprises accurate tokens. Additionally, the Pseudogene Correction Rate is expected to be greater than 25%, corresponding to the probability of randomly selecting one base out of four.

**Dataset Construction**: Pseudogene sequences were extracted from the human reference genome (GRCh38.p14) based on the GENCODE v46 annotation (Frankish et al., 2023). To identify paired pseudogenes and protein-coding genes, these pseudogene sequences were compared to a reference set of protein-coding gene translations (GENCODE v46) using a DIAMOND (Buchfink et al., 2021) BLASTX-like homology search in ultrasensitive mode, with an e-value threshold of $1 \times 10^{-10}$. The matches were filtered using an 80% identity cutoff. Each filtered pseudogene sequence was then aligned to its corresponding protein-coding gene using a nucleotide NCBI blastn search (Camacho et al., 2009). Only pseudogene–protein-coding gene pairs with a sequence length difference of $\leq$ 200 bp were retained to ensure high-quality pairings. This length constraint was implemented to prevent bias in subsequent model predictions. Finally, Smith-Waterman alignments, employing the BLASTN scoring matrix, were performed to record insertions, deletions, and mismatches in pseudogenes relative to their paired protein-coding genes. None of the samples include subsequences that are identical or partially identical to those present in the pre-trained dataset. The relevant data are accessible and can be validated through the publicly available dataset we have deposited on Zenodo: https://doi.org/10.5281/zenodo.15171943 (**Data Availability-Supplementary**).

We selected the samples whose mismatch rate does not exceed 30% and their sequence length does not exceed 4,096 (due to computational resource constraints), resulting in 160 samples (Dataset statistics are presented in **Supplementary Figure 13**).

The results are consistent with our hypothesis expectation for this task (**Supplementary Figure 14**). Firstly, the top-4 accuracy for all three experiments is 100%, indicating that the predictions do not deviate from A, T, C, or G to other tokens.

Secondly, the Gene Random Mask Recover Rate (0.5195) is significantly higher than the Pseudogene Keep Rate (0.3044), demonstrating that the LucaOne embedding is able to recognize the correct tokens in the gene sequence from problematic tokens.

Thirdly, the pseudogene correction rate is 0.3808, higher than 0.25, demonstrating its ability to rectify erroneous tokens within pseudogenes.

### 8 Task on ProtLoc with Nucleic Acid Embeddings

To further assess the efficacy and versatility of the LucaOne model in nucleic acid and protein-based tasks, an extended evaluation was performed using the ProtLoc dataset (details about **ProtLoc** task refer to **Methods. H**).

The protein sequences of the original task were mapped to nucleic acid sequences in the NCBI RefSeq database using their UniProt IDs, and the coding sequences (CDS) of the corresponding genes were extracted to construct a new dataset (refer to **Data Availability-Supplementary**). A few UniProt IDs lacked nucleic acid sequence mappings, resulting in a slight reduction in dataset size. Retaining the original training, validation, and testing splits, the final dataset included 8,005, 1,521, and 854 CDS sequences (see the ProLoc CDS-Gene and CDS-Prot datasets), respectively. The corresponding nucleic acid sequence embeddings were generated with LucaOne-5.6M (checkpoint 5.6M), and the model's performance was evaluated using the same framework as the original task.

From the results presented in **Supplementary Figure 15**, the performances of LucaOne and ESM2-3B on CDS-Prot were nearly identical. Among the two models based on nucleic acid data, LucaOne on CDS-Gene outperformed DNABert2 but remains slightly inferior to the two protein-based models. These findings suggested that embeddings derived from nucleic acid sequences are applicable to tasks related to their corresponding protein sequences, with LucaOne demonstrating relatively superior performance.

### 9 Task on SYN and NON-SYN Mutations Embeddings Distances

Firstly, we designed a task based on influenza virus HA sequence data to verify whether LucaOne can distinguish between synonymous and non-synonymous mutations in a zero-shot manner, more details in **Supplementary Figure 16-a**.

To examine the embedding distance offsets resulting from a larger number of synonymous and non-synonymous mutations, as well as to compare different models, we have prepared an additional test based on artificial mutations of real sequences (non-natural mutations). We collected coding gene sequences from ten species, including *Mus musculus*, *Chlamydia trachomatis D/UW-3/CX*, *Arabidopsis thaliana*, *Caenorhabditis elegans*, *Danio rerio*, *Saccharomyces cerevisiae S288C*, *Caulobacter vibrioides NA1000*, *Drosophila melanogaster*, *Coxiella burnetii RSA 493*, and *Homo sapiens*. Sequences shorter than 150 nucleotides were excluded, resulting in 7,045 genes.

Ten codons were randomly selected from each original sequence for single-nucleotide mutations. Synonymous and non-synonymous mutations were then performed at these sites, generating 20 mutated sequences per original sequence (10 synonymous and 10 non-synonymous mutations with mutation counts from 1 to 10).

These sequences were embedded into matrices using LucaOne or DNABert2, which were then transformed to vectors by max pooling (2,560 dimensions for LucaOne, 768 dimensions for DNABert2). Euclidean distances between original and mutated sequences (both non-synonymous and synonymous) were calculated, showing synonymous mutations caused less significant embedding changes (**Supplementary Figure 16-b, 16-c, and 16-d**).

LucaOne and DNABert2 were compared regarding their ability to distinguish synonymous and non-synonymous mutations across 10 species. Both models could discriminate between mutation types, but LucaOne demonstrated greater competence, particularly as mutation counts increased. All datasets are available; please see Section - **Data Availability-Supplementary**.

### 10 Task on Nearest Neighbors Analysis for Urochordates Species

This task is to analyze the mutual distances of gene sets from different species in the LucaOne embedding space. To expand species coverage, a more comprehensive dataset was constructed from reference and representative genomes of Eukaryota in NCBI RefSeq. After excluding unclassified classes, species containing over 500 genes were selected, and genes with nucleic acid sequences shorter than 2,500 bp were filtered for analysis. One species was randomly chosen from each class (prioritizing reference genomes), with 100 genes sampled per species. The final dataset encompassed 31 phyla, 94 classes, and 94 species plus three species from previous tasks: *Ciona intestinalis*, *Styela clava*, and *Oikopleura dioica*, in total 97 species.

The nucleic acid sequences of 100 genes selected from these species were embedded using LucaOne, and the Euclidean distances between different genes were calculated based on the $[CLS]$ vectors of each gene sequence embedding. The average of the pairwise embedding distances between all genes of the two species was used as the distance between the species (**Eq. (18)**).

$$Dist_{Sp_m Sp_n} = \frac{1}{|Sp_m| \times |Sp_n|} \sum_{i \in Sp_m,} \sum_{j \in Sp_n} dist(e_i, e_j) \tag{18}$$

where, $Sp_n$ is the sequences set of species $n$, $|Sp_n|$ is the size of $Sp_n$, $e_i$ is the embedding vector of sequence $i$, and $dist$ is the Euclidean distance function.

The list of all 97 species was presented in **Supplementary Table Sheet - Species to *Ciona intestinalis***, and also the distance between *Ciona intestinalis* and all other species. The top 5 nearest species among the 96 are *Styela clava*, *Dendronephthya gigantea*, *Corticium candelabrum*, *Danio rerio*, and *Saccoglossus kowalevskii*. All datasets are available; please see Section - **Data Availability-Supplementary**.

| Data Type/Source | Seq Count | Seq Length (Min/Max/Mean/Median) | Seq Count in which Annotation Exist |
|---|---|---|---|
| DNA/Refseq | 1,181,133,873 | 3/1,280/987/1,280 | Genome Region Type: 1,124,090,054 |
| | | | Order-level Taxonomy: 1,180,785,987 |
| RNA/Refseq | 136,311,178 | 3/1,280/1,035/1,280 | Genome Region Type: 106,918,452 |
| | | | Order-level Taxonomy: 135,933,074 |
| Protein/Uniref50 | 62,150,523 | 13/45,356/286/194 | Order-level Taxonomy: 58,537,832 |
| | | | Keyword: 35,188,544 |
| | | | Site: 3,341,857 |
| | | | Domain: 21,981,013 |
| | | | Homology: 24,246,265 |
| | | | Structure: 190,861 |
| Protein/UniProt | 252,170,925 | 3/45,356/352/281 | Order-level Taxonomy: 242,706,004 |
| | | | Keyword: 192,226,173 |
| | | | Site: 37,449,976 |
| | | | Domain: 141,276,963 |
| | | | Homology: 156,909,761 |
| | | | Structure: 569,034 |
| Protein/ColabFoldDB | 208,966,064 | 4/36,993/182/147 | Order-level Taxonomy: 5,389,742 |
| | | | Keyword: 3,117,716 |
| | | | Site: 343,344 |
| | | | Domain: 1,972,216 |
| | | | Homology: 2,136,440 |
| | | | Structure: 6,715 |

**Supplementary Figure 1**: **The statistics on pre-training data.** The inclusion of Uniref50, a subset of UniProt, enhances the learning of representative sequences. The explanation of terms (Genome Region Type, Order-level Taxonomy, Keyword, Site, Domain, Homology, and Structure) is in **Methods C.**

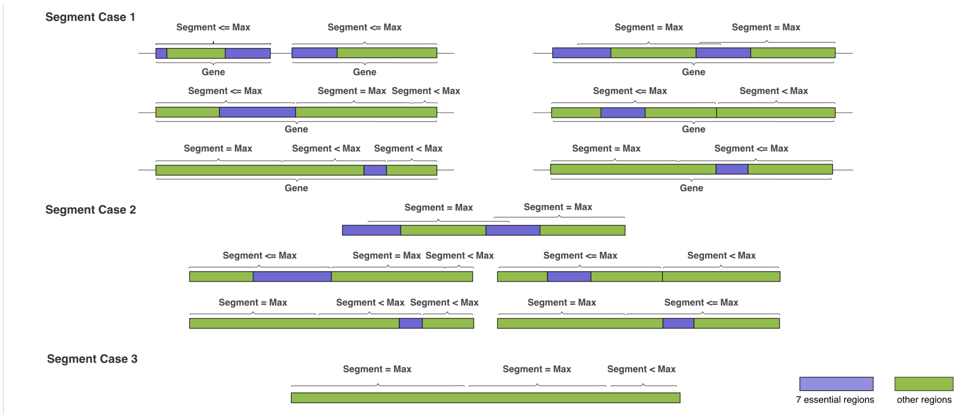

**Supplementary Figure 2**: **Three cases of segmentation.** 7 essential regions include: ’CDS’, ’regulatory’, ’tRNA’, ’ncRNA’, ’rRNA’, ’miscRNA’, and ’tmRNA’. Other regions include the genome region types not mentioned above, such as ’intron’. Each nucleic acid sequence was segmented according to a given maximum length. The fragmentation included the following three scenarios. **Case 1**: Only the genome regions were selected in the sequence for processing. If the length of consecutive genome regions did not exceed the maximum, merge them. It was segmented for genome regions exceeding the maximum length, and the fragment position should be ensured as far as possible in non-7 important regions (excluding intron regions). If a fragment was less than the maximum length, the fragment was expanded left and right simultaneously. **Case 2**: The entire sequence without genome regions was processed. If the length exceeded the maximum, it was fragmented in the same way as in **Case 1**. **Case 3**: For sequences without genome regions and 8 essential regions, the entire sequence was fragmented at no more than the specified maximum length.

| Task Level | Task | Task Type | Label Size | Description |
| --- | --- | --- | --- | --- |
| Token Level | Gene Mask | Multi-Class | 39 | Nucleotide mask |
| | Prot Mask | Multi-Class | 39 | Amino acid mask |
| Span Level | Genome Region Types | Multi-Class | 8 | Selected genome region types |
| | Prot Site | Multi-Class | 946 | Site region |
| | Prot Homo | Multi-Class | 3,443 | Homology region |
| | Prot Domain | Multi-Class | 13,717 | Domain region |
| Seq Level | Gene Taxonomy | Multi-Class | 735 | Order-level taxonomy |
| | Prot Taxonomy | Multi-Class | 2,196 | Order-level taxonomy |
| | Prot Keyword | Multi-Label | 1,179 | Protein keyword |
| Structure Level | Prot Structure | Regression | - | Amino-acid level position |

**Supplementary Figure 3**: **The pre-training tasks.** Details of the four levels of pretraining tasks are as follows: task name, task type, label size for classification task, and task description. The detailed description of all pre-training tasks is in **Methods D**.

| Task | Input | Method | Accuracy | F1-Score | PR-AUC | SRCC |
|---|---|---|---|---|---|---|
| CentralDogma | DNA-Protein | Transformer | 0.6662 | 0.0015 | 0.3372 | - |
| | | DNABert2 + ESM2-3B | 0.7309 | 0.5689 | 0.6065 | - |
| | | LucaOne (Gene/Prot) | 0.8048 | 0.6804 | 0.7750 | - |
| | | LucaOne | **0.8453** | **0.7392** | **0.8617** | - |
| SpeciesTax | DNA | BERTax★ [57] | - | - | - | - |
| | | DNABert2 | 0.696 | 0.6822 | 0.7645 | - |
| | | LucaOne | **0.750** | **0.7345** | **0.7927** | - |
| GenusTax | DNA | BERTax★ [57] | 0.124 | - | - | - |
| | | DNABert2 | 0.767 | 0.7562 | 0.8274 | - |
| | | LucaOne | **0.817** | **0.8069** | **0.8633** | - |
| SupKTax | DNA | BERTax★ [57] | 0.816 | - | - | - |
| | | DNABert2 | 0.848 | 0.8464 | 0.8747 | - |
| | | LucaOne | **0.947** | **0.9467** | **0.9591** | - |
| ProtLoc | Protein | DeepLocPro▲ [26] | 0.92 | - | - | - |
| | | ESM2-3B | **0.9496** | 0.9375 | 0.9659 | - |
| | | LucaOne | 0.9452 | **0.9378** | **0.9692** | - |
| ProtStab | Protein | TAPE▲ [51] | - | - | - | 0.73 |
| | | ESM2-3B | - | - | - | 0.7556 |
| | | LucaOne | - | - | - | **0.7718** |
| ncRNAFam | RNA | RNAGCN▲ [53] | 0.90 | - | - | - |
| | | DNABert2 | 0.9610 | 0.8444 | 0.9295 | - |
| | | LucaOne | **0.9864** | **0.9117** | **0.9622** | - |
| InfA | RNA-RNA | PREDAC● [54] | 0.9010 | 0.9279 | 0.9093 | - |
| | | DNABert2 | 0.9966 | 0.9941 | **1.0** | - |
| | | LucaOne | **1.0** | **1.0** | **1.0** | - |
| PPI | Protein-Protein | DeepPPI▲ [55] | 0.9719 | - | - | - |
| | | ESM2-3B | 0.9764 | 0.9790 | **0.9937** | - |
| | | LucaOne | **0.9774** | **0.9799** | 0.9922 | - |
| ncRPI | RNA-Protein | ncRPI-LGAT▲ [27] | **0.966** | - | - | - |
| | | DNABert2 + ESM2-3B | 0.9460 | 0.9450 | 0.9726 | - |
| | | LucaOne | 0.9479 | **0.9470** | **0.9781** | - |

**Supplementary Figure 4**: **More metrics of the testing set on downstream validation tasks.** The **macro** average strategy was adopted for F1-Score and PR-AUC of all multi-class tasks. The top right ★ indicates inference using the trained method, the top right ▲ indicates direct use of the results in its paper, and the top right ● indicates repetition using its method and higher than the results in the paper.

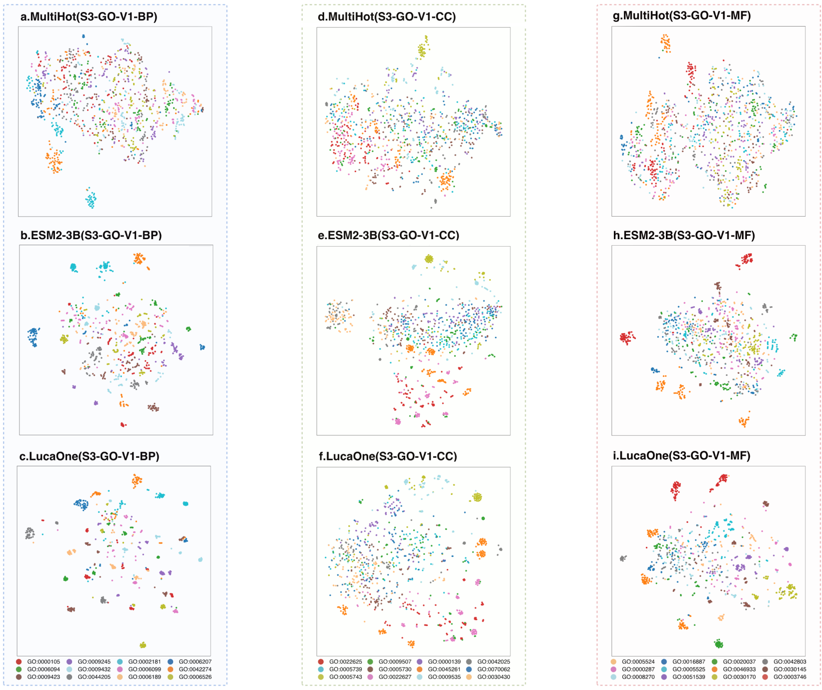

**Supplementary Figure 5**: **Comparative visualization of embedding methods on GO V1 dataset.** t-SNE projections of MultiHot encoding, ESM2-3B embeddings, and LucaOne embeddings across three Gene Ontology (GO) categories (biological processes [BP], cellular components [CC], molecular functions [MF]) for the S3-GO-V1 dataset. LucaOne consistently demonstrates clearer separation and clustering patterns compared to ESM2-3B and MultiHot, particularly in the BP and MF categories, highlighting its superior capability for GO annotation tasks (Comparison of clustering metrics in **Supplementary Figure 7**).

**S3-GO-V1-BP: GO:0000105**: L-histidine biosynthetic process, **GO:0009245**: lipid A biosynthetic process, **GO:0002181**: cytoplasmic translation, **GO:0006207**: 'de novo' pyrimidine nucleobase biosynthetic process, **GO:0006094**: gluconeogenesis, **GO:0009432**: SOS response, **GO:0006099**: tricarboxylic acid cycle, **GO:0042274**: ribosomal small subunit biogenesis, **GO:0009423**: chorismate biosynthetic process, **GO:0044205**: 'de novo' UMP biosynthetic process, **GO:0006189**: 'de novo' IMP biosynthetic process, **GO:0006526**: L-arginine biosynthetic process;

**S3-GO-V1-CC: GO:0022625**: cytosolic large ribosomal subunit, **GO:0009507**: chloroplast, **GO:0000139**: Golgi membrane, **GO:0042025**: host cell nucleus, **GO:0005739**: mitochondrion, **GO:0005730**: nucleolus, **GO:0045261**: proton-transporting ATP synthase complex, catalytic core F(1), **GO:0070062**: extracellular exosome, **GO:0005743**: mitochondrial inner membrane, **GO:0022627**: cytosolic small ribosomal subunit, **GO:0009535**: chloroplast thylakoid membrane, **GO:0030430**: host cell cytoplasm;

**S3-GO-V1-MF: GO:0005524**: ATP binding, **GO:0016887**: ATP hydrolysis activity, **GO:0020037**: heme binding, **GO:0042803**: protein homodimerization activity, **GO:0000287**: magnesium ion binding, **GO:0005525**: GTP binding, **GO:0046933**: proton-transporting ATP synthase activity, rotational mechanism, **GO:0030145**: manganese ion binding, **GO:0008270**: zinc ion binding, **GO:0051539**: 4 iron, 4 sulfur cluster binding, **GO:0030170**: pyridoxal phosphate binding, **GO:0003746**: translation elongation factor activity.

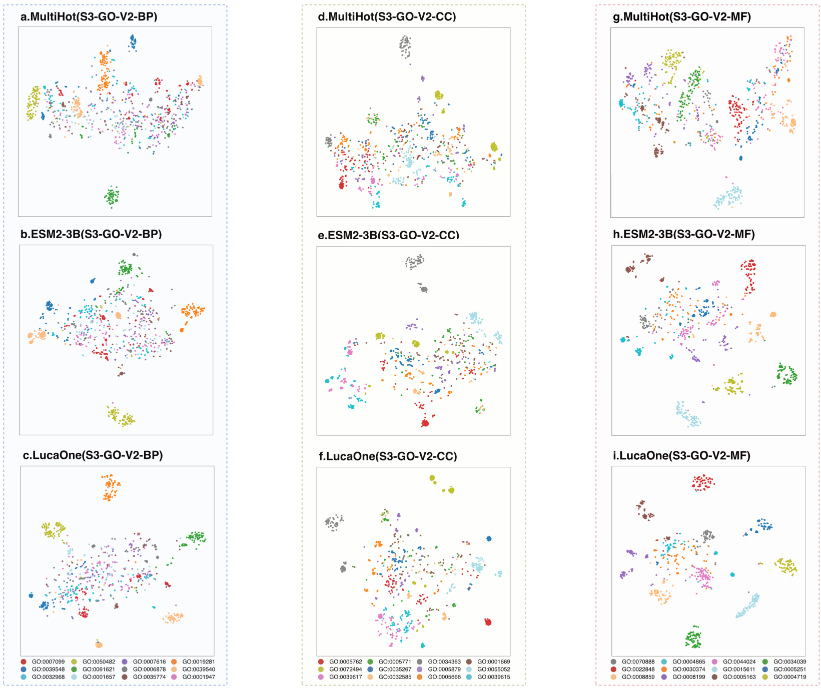

**Supplementary Figure 6**: **Comparative visualization of embedding methods on GO V2 datasets.** t-SNE projections of MultiHot encoding, ESM2-3B embeddings, and LucaOne embeddings across three Gene Ontology (GO) categories (biological processes [BP], cellular components [CC], molecular functions [MF]) for the S3-GO-V2 dataset. The specific numerical comparisons are referred to in **Supplementary Figure 7**.

**S3-GO-V2-BP: GO:0007099**: centriole replication, **GO:0050482**: arachidonate secretion, **GO:0007616**: long-term memory, **GO:0019281**: L-methionine biosynthetic process from homoserine via O-succinyl-L-homoserine and cystathionine, **GO:0039548**: symbiont-mediated suppression of host cytoplasmic pattern recognition receptor signaling pathway via inhibition of IRF3 activity, **GO:0061621**: canonical glycolysis, **GO:0006878**: intracellular copper ion homeostasis, **GO:0039540**: symbiont-mediated suppression of host cytoplasmic pattern recognition receptor signaling pathway via inhibition of RIG-I activity, **GO:0032968**: positive regulation of transcription elongation by RNA polymerase II, **GO:0001657**: ureteric bud development, **GO:0035774**: positive regulation of insulin secretion involved in cellular response to glucose stimulus, **GO:0001947**: heart looping;

**S3-GO-V2-CC: GO:0005762**: mitochondrial large ribosomal subunit, **GO:0005771**: multivesicular body, **GO:0034363**: intermediate-density lipoprotein particle, **GO:0001669**: acrosomal vesicle, **GO:0072494**: host multivesicular body, **GO:0035267**: NuA4 histone acetyltransferase complex, **GO:0005879**: axonemal microtubule, **GO:0055052**: ATP-binding cassette (ABC) transporter complex, substrate-binding subunit-containing, **GO:0039617**: T=3 icosahedral viral capsid, **GO:0032585**: multivesicular body membrane, **GO:0005666**: RNA polymerase III complex, **GO:0039615**: T=1 icosahedral viral capsid;

**S3-GO-V2-MF: GO:0070888**: E-box binding, **GO:0004865**: protein serine/threonine phosphatase inhibitor activity, **GO:0044024**: histone H2AS1 kinase activity, **GO:0034039**: 8-oxo-7,8-dihydroguanine DNA N-glycosylase activity, **GO:0022848**: acetylcholine-gated monoatomic cation-selective channel activity, **GO:0030374**: nuclear receptor coactivator activity, **GO:0015611**: ABC-type D-ribose transporter activity, **GO:0005251**: delayed rectifier potassium channel activity, **GO:0008859**: exoribonuclease II activity, **GO:0008199**: ferric iron binding, **GO:0005163**: nerve growth factor receptor binding, **GO:0004719**: protein-L-isoaspartate (D-aspartate) O-methyltransferase activity.

| DataSet | Embedding | ARI | AMI | HS | CS | V-measure | FMI |
|---|---|---|---|---|---|---|---|
| S3-GO-V1-BP | Multi-OneHot | 0.0942 | 0.1920 | 0.1934 | 0.2315 | 0.2107 | 0.2004 |
| | ESM2-3B | 0.1079 | 0.2582 | 0.2596 | 0.2928 | 0.2752 | 0.2027 |
| | LucaOne | **0.1535** | **0.3779** | **0.3707** | **0.4159** | **0.3920** | **0.2474** |
| S3-GO-V1-CC | Multi-OneHot | 0.0707 | 0.1510 | 0.1557 | 0.1911 | 0.1716 | 0.1843 |
| | ESM2-3B | 0.1384 | 0.2861 | 0.2854 | 0.3218 | 0.3025 | 0.2303 |
| | LucaOne | **0.1507** | **0.3220** | **0.3245** | **0.3505** | **0.3370** | **0.2353** |
| S3-GO-V1-MF | Multi-OneHot | 0.0519 | 0.1082 | 0.1112 | 0.1530 | 0.1288 | 0.1791 |
| | ESM2-3B | 0.1008 | 0.2476 | 0.2538 | 0.2758 | 0.2643 | 0.1899 |
| | LucaOne | **0.1777** | **0.4000** | **0.3935** | **0.4356** | **0.4135** | **0.2672** |
| S3-GO-V2-BP | Multi-OneHot | 0.1268 | 0.2324 | 0.2195 | 0.2919 | 0.2506 | 0.2488 |
| | ESM2-3B | 0.2977 | 0.4464 | 0.4500 | 0.4669 | 0.4583 | 0.3622 |
| | LucaOne | **0.3394** | **0.4983** | **0.4967** | **0.5224** | **0.5092** | **0.4029** |
| S3-GO-V2-CC | Multi-OneHot | 0.0970 | 0.2259 | 0.2102 | 0.2926 | 0.2446 | 0.2304 |
| | ESM2-3B | 0.2872 | 0.4577 | 0.4555 | 0.4846 | 0.4696 | 0.3564 |
| | LucaOne | **0.3385** | **0.5142** | **0.4993** | **0.5541** | **0.5253** | **0.4110** |
| S3-GO-V2-MF | Multi-OneHot | 0.2256 | 0.3900 | 0.3609 | 0.4603 | 0.4046 | 0.3360 |
| | ESM2-3B | 0.4699 | 0.6524 | 0.6450 | 0.6756 | 0.6599 | 0.5207 |
| | LucaOne | **0.6608** | **0.7796** | **0.7727** | **0.7962** | **0.7843** | **0.6919** |

**Supplementary Figure 7**: **Clustering metrics (using K-Means++).** The figure evaluates Multi-OneHot, ESM2-3B, and LucaOne embeddings across six clustering metrics (ARI, AMI, HS, CS, V-measure, FMI) for S3-GO-V1 and V2 datasets spanning biological processes [BP], cellular components [CC], and molecular functions [MF]. LucaOne consistently outperforms baseline methods in all metrics, with particularly notable gains in the S3-GO-V2-MF subset. The V2 datasets were constructed by selecting GO terms with minimal semantic overlap to mitigate inherent term correlations in V1, enabling clearer discriminative analysis.

| Method | Embedding | Person | | Spearman | |
|---|---|---|---|---|---|
| | | Correlation Coefficient ↑ | P-value ↓ | Correlation Coefficient ↑ | P-value ↓ |
| Lin Similarity V.S. Centroids Cosine Similarity | ESM2-3B | 0.6076 | 6.75e-16 | 0.6420 | 4.28e-18 |
| | LucaOne | **0.7143** | **9.11e-24** | **0.7089** | **2.77e-23** |
| Lin Similarity V.S. Mean Pairwise Cosine Similarity | ESM2-3B | 0.5329 | 6.16e-12 | 0.2499 | 2.52e-03 |
| | LucaOne | **0.7481** | **4.58e-27** | **0.7356** | **8.60e-26** |
| Lin Similarity V.S. Median Pairwise Cosine Similarity | ESM2-3B | 0.5931 | 4.79e-15 | 0.3910 | 1.26e-06 |
| | LucaOne | **0.7312** | **2.37e-25** | **0.6646** | **1.05e-19** |

**Supplementary Figure 8**: **Correlation analysis between GO term similarity and sequence embedding similarity.** This table presents the Pearson and Spearman correlation coefficients, along with their corresponding p-values, comparing Lin similarity of GO terms with three different sequence embedding-based similarity measures: **(1)** centroids cosine similarity, **(2)** mean pairwise cosine similarity and **(3)** median pairwise cosine similarity. The analysis was performed separately for ESM2-3B and LucaOne embeddings.

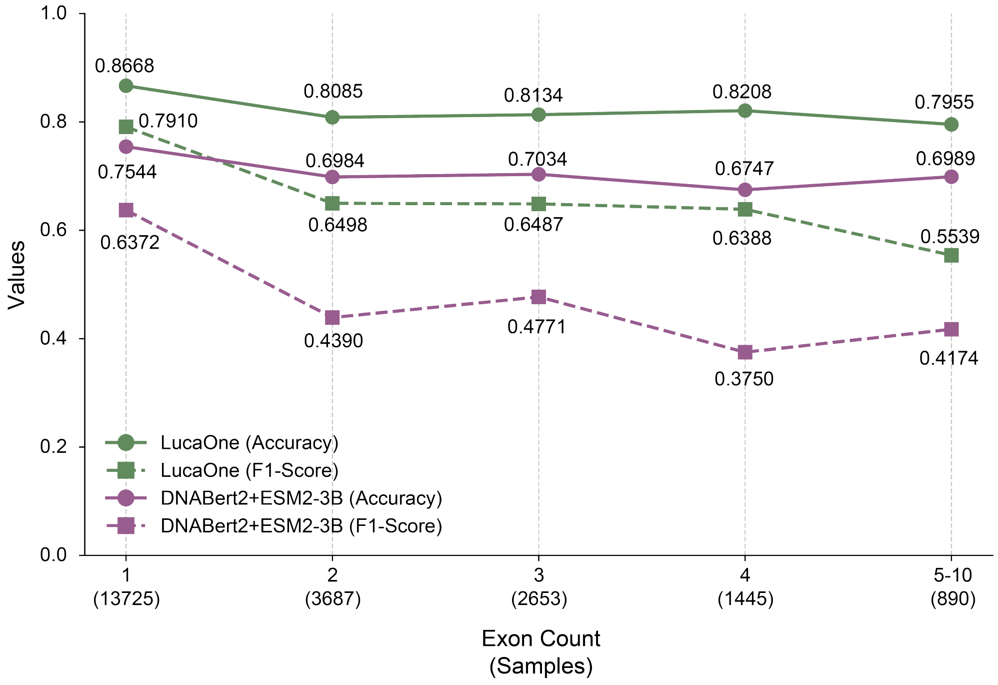

**Supplementary Figure 9**: **Comparative performance analysis across different exon count subsets.** LucaOne consistently outperformed DNABert2 + ESM2-3B in both Accuracy and F1-Score across all exon count subsets. Data Details see **Supplementary Data Set - Exon Performance by sp.**

| DataSet train: valid: test | Embedding | Network | Accuracy (test) | F1-Score (test) | AUC (test) | PR-AUC (test) |
|---|---|---|---|---|---|---|
| 4: 3: 25 | DNABert2 + ESM2-3B | Pooling + FC | 0.5486 | 0.3248 | 0.4944 | 0.3264 |
| | LucaOne | | **0.5571** | **0.3367** | **0.5017** | **0.3290** |
| 8: 1: 1 | DNABert2 + ESM2-3B | Pooling + FC | 0.7173 | 0.6085 | 0.7745 | 0.6445 |
| | LucaOne | | **0.7587** | **0.6577** | **0.8167** | **0.6865** |

**Supplementary Figure 10**: **Central Dogma tasks cross-species homologous gene pairs.** Across-species homologous gene pairs of the Central Dogma task. The dataset was constructed by pairing nucleic acid and protein sequences of the same gene across different species. Here is the comparative analysis of model performance across various embedding models and data partition ratios.

| Superkingdom | Species | Original Pairs Count | Sampling Pairs Count |
|---|---|---|---|
| Viruses | Human immunodeficiency virus | 1 | / |
| | Tequatrovirus | 7 | / |
| Bacteria | Escherichia coli | 2 | / |
| | Salmonella enterica | 2 | / |
| | Staphylococcus elegans | 2 | / |
| | Coxiella burnetii | 44 | 44 |
| | Chlamydia trachomatis | 81 | 81 |
| | Caulobacter vibrioides | 263 | 200 |
| Eukaryota | Homo sapiens | 31 | 31 |
| | Mus musculus | 55 | 55 |
| | Danio rerio | 73 | 73 |
| | Saccharomyces cerevisiae | 541 | 200 |
| | Drosophila melanogaster | 1,051 | 200 |
| | Arabidopsis thaliana | 1,557 | 200 |
| | Caenorhabditis eleganss | 4,108 | 200 |
| Total | / | 7,818 | 1,284 |

**Supplementary Figure 11**: **Dataset composition for cross-species gene-protein pair analysis.** The table summarizes the construction of a filtered dataset comprising 1,284 gene-protein pairs across 10 species from three superkingdoms (Viruses, Bacteria, and Eukaryota). Species with over 200 genes were subsampled to 200 pairs, while others retained all qualifying pairs. Sequences were excluded if they aligned with the trained sequences of LucaOne-5.6M (checkpoint 5.6M) via NCBI blastn (nucleic acids) or Diamond blastp (proteins).

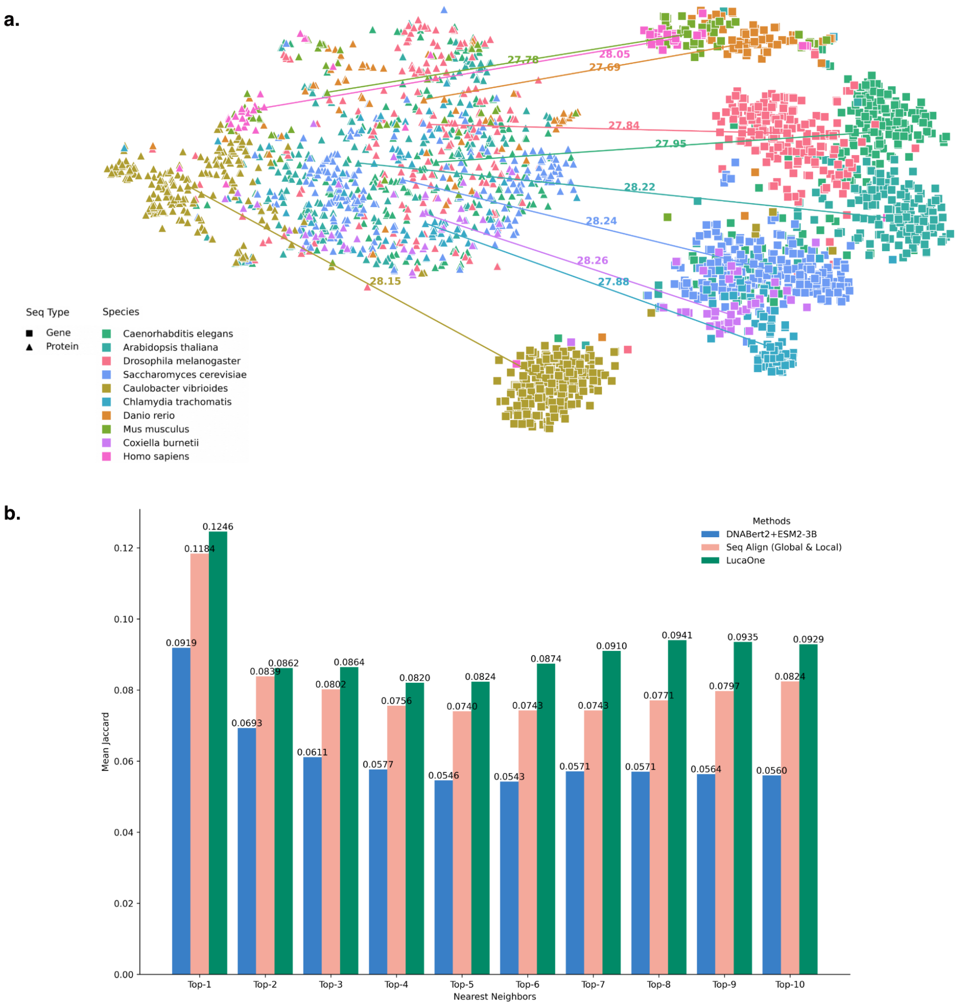

**Supplementary Figure 12**: **t-SNE and Jaccard Index analysis for 1,284 genes. a.** t-SNE for 1,284 pairs of nucleic acid and protein sequences for the same gene. Nucleic acid and protein sequences were consistently coloured based on their corresponding gene, with proteins represented by triangles and nucleic acids by squares. **b.** Comparative analysis of mean Jaccard scores across different methods for Top 1∼10 nearest neighbors. This figure presents a comparative analysis of the mean Jaccard similarity coefficients for the Top 1∼10 nearest neighbors, evaluated across three distinct methodologies: DNABert2 + ESM2-3B embedding, sequence alignment (both global and local alignment), and LucaOne embedding. The results demonstrate that LucaOne consistently outperforms the other methods across all Top-K evaluations, with DNABert2 + ESM2-3B showing the lowest performance.

| Count | | | |
|---|---|---|---|
| Total Sample Count | Total Token Count | Total Mask Token Count | |
| 160 | 131,034 | 11,009 | |
| Sequence Len | | | |
| Mean | Median | Min | Max |
| 818.96 | 681 | 180 | 3,399 |
| Mask Token Count | | | |
| Mean | Median | Min | Max |
| 68.81 | 53 | 1 | 551 |
| Mask Token Probability | | | |
| Mean | Median | Min | Max |
| 0.09167 | 0.086 | 0.0014 | 0.2896 |

**Supplementary Figure 13**: **Dataset statistics for pseudogene correction task.** The dataset details for the pseudogene correction task are as follows: number of samples, total token count, masked token count, sequence length statistics, masked token count statistics, and masked token probability statistics.

| Task | Accuracy | Top2-Accuracy | Top4-Accuracy |
|---|---|---|---|
| Gene Random Mask Recover | 0.5195 | 0.7764 | 1.0 |
| Pseudogene Keep | 0.3044 | 0.5852 | 1.0 |
| Pseudogene Correction | 0.3808 | 0.6601 | 1.0 |

**Supplementary Figure 14**: **Results for pseudogene correction task.** The Gene Random Mask Recover Rate (0.5195) is higher than the Pseudogene Keep Rate (0.3044), demonstrating that the LucaOne embedding is able to recognize the correct tokens in the gene sequence from problematic tokens. The pseudogene correction rate is 0.3808, higher than 0.25, demonstrating its ability to rectify erroneous tokens within pseudogenes.

**a.**

| ProtLoc | Original Dataset | | Corresponding CDS-Gene Dataset | | Corresponding CDS-Prot Dataset | |
|---|---|---|---|---|---|---|
| Dataset | Seq Count | Proportion | Seq Count | Proportion | Seq Count | Proportion |
| Training | 9915 | 76.05% | 8005 | 77.12% | 8005 | 77.12% |
| Validation | 1991 | 15.27% | 1521 | 14.65% | 1521 | 14.65% |
| Testing | 1131 | 8.68% | 854 | 8.23% | 854 | 8.23% |
| Total | 13037 | 100% | 10380 | 100% | 10380 | 100% |

**b.**

| Task | Dataset | Embedding | Network | Accuracy | F1-Score | PR-AUC |
|---|---|---|---|---|---|---|
| ProtLoc | Original | ESM2-3B | Attention Pooling + FC | **0.9496** | 0.9375 | 0.9659 |
| | | LucaOne | | 0.9452 | **0.9378** | **0.9692** |
| | CDS-Gene | DNABert2 | Attention Pooling + FC | 0.8993 | 0.9005 | 0.9329 |
| | | LucaOne | | **0.9379** | **0.9226** | **0.9494** |
| | CDS-Prot | ESM2-3B | Attention Pooling + FC | **0.9520** | 0.9434 | 0.9676 |
| | | LucaOne | | 0.9496 | **0.9438** | **0.9690** |

**Supplementary Figure 15**: **Task on ProLoc with nucleic acid embeddings: the dataset statistics and performance. a.** the dataset statistics for original ProtLoc and corresponding CDS-Gene and CDS-Gene-Prot. **b.** The performance comparison between LucaOne and DNABert2/ESM2-3B on original and CDS-Gene/CDS-Prot datasets. The colored background represents two newly generated datasets based on the original dataset.

**a.**

**1.**

| Amino Acid Mutation | Nucleotide Mutation | Pair Count |
|---|---|---|
| 1 | 1 | 209 |
| 2 | 2 | 705 |
| 3 | 3 | 354 |

**2.**

| Number of Mutation | Average Euclidean Embeddings Distance (Synonymous) | Average Euclidean Embeddings Distance (Non-Synonymous) | P-Value (t-test) |
|---|---|---|---|
| 1 | 0.06889 | 0.09829 | 1.82408e-10 |
| 2 | 0.11437 | 0.15470 | 8.69506e-34 |
| 3 | 0.14094 | 0.18606 | 1.84772e-18 |

**b.**

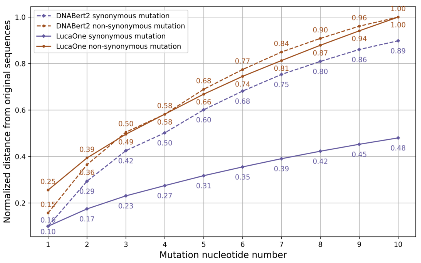

**c.**

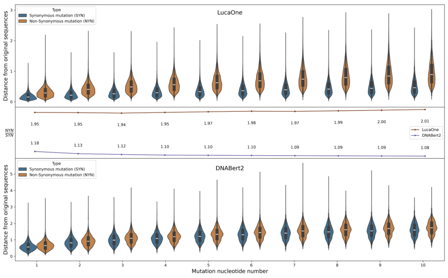

**d.**

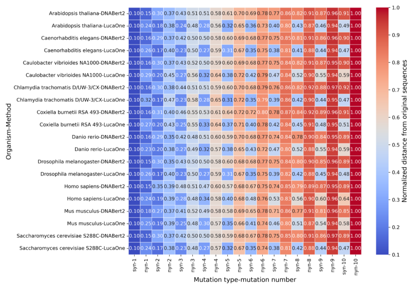

**Supplementary Figure 16**: **Task on codon degeneracy: data statistics and embeddings distance. a.** The count of sequence pairs with one, two, or three amino acid mutations has been assembled, with each mutation resulting from a single nucleotide change, and the differences of the embedded max pooling vectors between the original sequence and sequences containing 1, 2, or 3 non-synonymous mutations or synonymous mutations. Dataset refers to **Methods F. LucaOne Embeddings Level Analysis - Task on Codon Degeneracy**. **b.** The line plot of normalized distances between mutated and original sequences. The distances are normalized as the tips. For each data point, we calculate the mean distance of all species. **c.** The violin plot of the original distance (unnormalized) and the line plot of fold change between non-synonymous and synonymous mutation distance. **d.** The Heatmap of normalized distances between mutated and original sequences. The distances are normalized as the tips. Dataset for **b.**, **c.**, and **d.** refer to **Supplementary Note 9: Task on SYN and NON-SYN Mutations Embeddings Distances.**

| Terms & Abbreviation | Definition |
|---|---|
| NLP | Natural Language Processing |
| SOTA | State of the Art |
| Transformer | a Popular Deep Learning Architecture |
| BERT | Bidirectional Encoder Representations from Transformers |
| RoPE | Rotary Positional Encoding |
| CDS | Coding Sequence |
| Intron | a Segment of DNA not Translated into Protein |
| tRNA | Transfer RNA |
| ncRNA | Non-Coding RNA |
| rRNA | Ribosomal RNA |
| miscRNA | Miscellaneous RNA |
| tmRNA | Transfer-Messenger RNA |
| Regulatory | the Regulation of Gene Expression |
| MMSeqs | Many-against-Many Sequence Searching |
| OneHot | Convert a Unique Category Value into a Binary Vector |
| Multi-OneHot | Convert Multiple Categories into a Binary Matrix based on OneHot |
| Taxonomy | a Hierarchical Classification System in Biology |
| Site | Protein Site Region |
| Homology | a Common Evolutionary Origin in Proteins |
| Domain | a Distinct Functional, Structural, or Sequence Unit |
| Keyword | Keywords for Protein Annotation |
| Structure | Three-Dimensional Arrangements of Atoms within a Protein |
| $C_\alpha$-*atom* | the Carbon that is Next to a Functional Group |
| RefSeq | NCBI Reference Sequence Database |
| UniProt | the Universal Protein Knowledgebase |
| UniRef | Comprehensive and Non-redundant UniProt Reference Clusters |
| ColabFoldDB | MMseqs2 Expandable Profile Databases of Proteins |
| InterPro | Classification of Protein Families |
| RCSB-PDB | Protein Data Bank |
| AlphaFold2 | AlphaFold2 Protein Structure Database |
| CAMI2 | Critical Assessment of Metagenome Interpretation II |
| Pfam | a Database with Large Collection of Protein Families |
| GO | Gene Ontology |
| ESM2-3B | a Protein Language Model with 3B Parameters |
| DNABert2 | a Transformer-based Genome Foundation Model |
| Pooling | Downsampling Operation that Reduces the Dimensionality of the Feature Map |
| $[CLS]$ Vector | Token $[CLS]$ embedding Vector |
| Mean Pooling | Each Feature Retains the Mean Value in Pooling |
| Max Pooling | Each Feature Retains the Maximum Value in Pooling |
| Value-Level Attention Pooling | Using the Attention Mechanism in Pooling |
| FC | Fully Connected Layer |
| Warm-up | Gradually Increasing the Learning Rate During the Initial Stages of Training |
| TP/TN/FP/FN | Four Metrics in the Confusion Matrix |
| Accuracy | Accuracy Classification Score |
| F1-Score | the Harmonic Mean of the Precision and Recall |
| AUC | the Area under the ROC-Curve, |
| PR-AUC | the Area Under the Precision-Recall Curve |
| SRCC | Spearman's Rank Correlation Coefficient |
| t-SNE | t-Distributed Stochastic Neighbor Embedding |
| ARI | Adjusted Rand Index Score |
| AMI | Adjusted Mutual Information Score |
| HS | the Homogeneity Score |
| CS | the Completeness Score |
| V-measure | the Harmonic Mean between Homogeneity and Completeness |
| FMI | the Fowlkes-Mallows Index Score |
| Jaccard Similarity | the Similarity between Two Sets |
| Lin Similarity | Node-based Semantic Similarity Measure |
| Pearson | Pearson Correlation Coefficient |
| Spearman | Spearman Correlation Coefficient |

**Supplementary Figure 17**: **Terms & Abbreviations definitions.** The full names and hyperlinks to details of the terms involved in the paper.
